# Functional and druggability analysis of the SARS-CoV-2 proteome

**DOI:** 10.1101/2020.08.21.261404

**Authors:** Claudio N. Cavasotto, Maximiliano Sánchez Lamas, Julián Maggini

## Abstract

The infectious coronavirus disease (COVID-19) pandemic, caused by the coronavirus SARS-CoV-2, appeared in December 2019 in Wuhan, China, and has spread worldwide. As of today, more than 22 million people have been infected, with almost 800,000 fatalities. With the purpose of contributing to the development of effective therapeutics, this work provides an overview of the viral machinery and functional role of each SARS-CoV-2 protein, and a thorough analysis of the structure and druggability assessment of the viral proteome. All structural, non-structural, and accessory proteins of SARS-CoV-2 have been studied, and whenever experimental structural data of SARS-CoV-2 proteins were not available, homology models were built based on solved SARS-CoV structures. Several potential allosteric or protein-protein interaction druggable sites on different viral targets were identified, knowledge that could be used to expand current drug discovery endeavors beyond the cysteine proteases and the polymerase complex. It is our hope that this study will support the efforts of the scientific community both in understanding the molecular determinants of this disease and in widening the repertoire of viral targets in the quest for repurposed or novel drugs against COVID-19.

## 1 Introduction

In the past decades, two highly pathogenic coronaviruses (CoVs), the severe acute respiratory syndrome coronavirus (SARS-CoV), and the Middle East respiratory syndrome coronavirus (MERS-CoV), triggered global epidemics in 2003 and 2012, respectively [1]. A new CoV infectious disease (COVID-19), caused by the new pathogenic SARS-CoV-2 appeared in December 2019 in Wuhan, China, spread rapidly worldwide, and was declared a pandemic on March 11^th^, 2020 by the World Health Organization (WHO). As this contribution is being written, there have been over 22 million infected cases around the world, with almost 800,000 fatalities. The scientific community has displayed a swift response to this pandemic, especially by investigating the molecular basis of this disease, which could help the quick development of effective antivirals and vaccines.

SARS-CoV-2 belongs to the β-coronavirus genus, like SARS-CoV and MERS-CoV. They are enveloped positive single-strain RNA (+ssRNA) viruses, with large genomes that give them great protein-coding capacity. Once the virus binds to the cell host receptor, their membranes fuse, and the viral genome is released into the cell cytoplasm. Once in the cytoplasm, the +ssRNA viral genome is translated to produce the viral proteins necessary to replicate the viral genome, and the structural proteins which assemble the new viral particles. The replicative proteins make several copies of the +ssRNA, and finally, the produced +ssRNA and structural proteins are assembled into new viral particles, which are then released from the host cell to restart the virus cycle [2].

Although SARS-CoV-2, SARS-CoV, and MERS-CoV belong to the same genus, SARS-CoV-2 seems to be associated with milder infections. Moreover, SARS and MERS were associated mainly with intrahospitalary and zoonotic transmission, whereas SARS-CoV-2 is much more widely spread within the community [3]. SARS-CoV-2 shares ∼94% sequence identity in replicative genes to SARS-CoV [4], while MERS is phylogenetically more distant. SARS-CoV and SARS-CoV-2 use the same host receptor, the angiotensin-converting enzyme 2 (ACE2), while MERS uses the dipeptidyl peptidase-4 (DPP4, also known as CD26) [5]. Clinically, SARS-CoV-2, SARS-CoV, and MERS-CoV have similar symptomology; they differ mainly in their fatality rates of 2.3%, 9.5%, and 34.4% respectively, and their basic reproductive numbers (R0) 2.0–2.5, 1.7– 1.9, and <1, respectively [6].

The typical clinical presentation of severe cases of COVID-19 is pneumonia with fever, cough, and dyspnea. The vast majority of patients have mild disease (around 80%), a lesser percentage (∼15%) have moderate disease (with dyspnea, hypoxia or pneumonia with over 50% lung parenchyma involvement), and even a smaller proportion (∼5%) of patients develop severe disease with respiratory failure, shock or multiorgan failure (5%) [7].

Although the factors that condition the progression of the disease in severe cases are not fully understood, evidence seems to show that the symptoms in patients with moderate to severe disease due to SARS-CoV-2 are related not only to viral proliferation, but also to two factors related to its pathogenesis: i) an exacerbated inflammatory response associated with increased concentrations of proinflammatory cytokines, such as tumor necrosis factor-α (TNF-α) and interleukins (IL), including IL-1 and IL-6 [8-11], and ii) abnormalities in coagulation and thrombosis, similar to a combination of mild disseminated intravascular coagulopathy and a localized pulmonary thrombotic microangiopathy, which could have a substantial impact on organ dysfunction in the more severely affected patients [12].

Although the optimal strategy to control this disease would be through vaccination capable of generating a long-lasting and protective immune response, there is an urgent need for the development of treatment for this disease, dictated by: i) the rapid spread of the virus; ii) the increasing number of infected patients with a moderate to severe pathology that must be treated worldwide; iii) the associated risk of death and depletion of the health systems, particularly in low-income countries. Moreover, since 20 to 30% of patients hospitalized for pneumonia require intensive care for mechanical ventilation [13,14], even a treatment to decrease the severity of the disease would be highly welcomed.

In a recent work, the SARS-CoV-2-human interactome was characterized, and +300 protein-protein interactions (PPIs) were identified. It was found that 66 of the interacting host factors could be modulated with 69 compounds, among these some approved drugs [15]. This study will certainly enhance the possibilities of drug discovery efforts targeting host cells receptors. Today, candidates include the antimalarial chloroquine and its derivatives [16], anti-inflammatory drugs and immunomodulators, and even anticoagulants and anti-fibrinolytic drugs [17].

In terms of antiviral strategies, while no specific therapeutics are available, approved or experimental drugs developed for other diseases are being tested in different clinical trials, in an effort to rapidly find accessible treatments with already established safety profiles, through drug repurposing strategies [18-23]. So far, most of these strategies are focused in targeting the two viral proteases, and the RNA-dependent RNA polymerase (RdRp) complex. However, protease inhibitors might bind to host proteases, thus resulting in cell toxicity and side effects [24]; the effect of nucleoside inhibitors targeting RdRp is decreased by the highly efficient SARS-CoV-2 proofreading machine, and even limitations in targeting the S protein have been pointed out [25]. Therefore, alternative therapeutic options to fight COVID-19 are badly needed.

In this context, characterization of the druggability of the SARS-CoV-2 proteome is of critical importance. In this work, we analyzed the functional role of each SARS-CoV-2 protein, and performed a thorough structural and druggability analysis of the viral proteome, using either experimental structural data or homology models, studying all structural, non-structural, and accessory proteins. Current drug discovery efforts on the proteases and the RdRp complex with experimental *in vitro* or *in vivo* validation are also briefly commented, mainly to complement our description of targetability. As described throughout this work, all viral proteins have important functions, and many of them could be a potential target for drug development, not only aiming to block virus entry or replication, but also to interfere with other viral aspects, such as genome stability (by inhibiting critical modifications such as capping and polyadenylation), viral particle assembly, immune evasion, and derived pathologies as inflammation and thrombosis. Besides the catalytic sites of several proteins, we found potential druggable allosteric and PPI sites throughout the whole proteome, including alternative sites within the proteases and the RdRp.

To find and assess the targetability of potential binding sites (including allosteric ones), we used FTMap [26-28] to identify binding hot spots on all available (or modeled) SARS-CoV-2 structures. FTMap samples through docking a library of a variety of small organic probes of different molecular properties on the protein. The poses of each probe are clustered, keeping the six lowest-energy clusters per probe; and then the probe clusters of all molecules are re-clustered into consensus sites (CSs) or hot spots. Hot spots are small regions within a binding site considered to strongly contribute to the ligand-binding free energy. This analysis was complemented by the use of ICM Pocket Finder [29], and by the detection of cryptic sites using CryptoSite [30], where cryptic sites are those usually “hidden” in unligated structures, but present only in ligand-bound structures. The strength of a hot spot (or consensus sites, CS) is related to the number of fragment-like probe clusters it contains. A strong CS and nearby hot spots define a potential binding site [26], which is considered druggable if its primary hot spot includes at least 16 probe clusters, and at least another hot spot within a certain threshold distance is present; these conditions would assure that a drug-like molecule might bind the receptor at least with micromolar affinity [26] (see Methods for a detailed description of this methodology). It is clear that this assessment of druggability does not necessarily mean that a ligand binding at that site will actually exert an observable biological effect [31].

We do hope our contribution will help in the development of fast and effective SARS-CoV-2-centered therapeutic options, which considering the high similarity among CoVs, might also be effective against related viruses. Moreover, since it is unlikely that SARS-CoV-2 will be the last CoV to threaten global health, those therapeutics might be instrumental to fight future epidemics.

## 2 The SARS-CoV-2 viral machinery

SARS-CoV-2 genetically clusters with the β-coronavirus genus, and is phylogenetically closely related to SARS-CoV. Its +ssRNA, of ∼30 kbs [32], is enclosed in a spherical lipidic bilayer membrane. About the first two-thirds of the viral RNA genome contain the open reading frames (ORFs) 1a and 1ab, which are translated into polyproteins (pp) 1a and 1ab, which contain the non-structural proteins (nsp’s). The remaining viral genome encode accessory proteins and four essential structural proteins [33]: the spike (S) receptor binding glycoprotein (ORF2); the nucleocapsid (N) protein (ORF9a); the membrane (M) protein (ORF5), a transmembrane (TM) protein involved in the interaction with N; and a small envelop (E) protein (ORF4), which participates in membrane stability [34] (Figure 1). The accessory proteins act co-opting host factors, shutting down host functions to redirect resources to viral replication, avoiding immune responses, or inducing pathogenicity [32].

**Figure 1.**
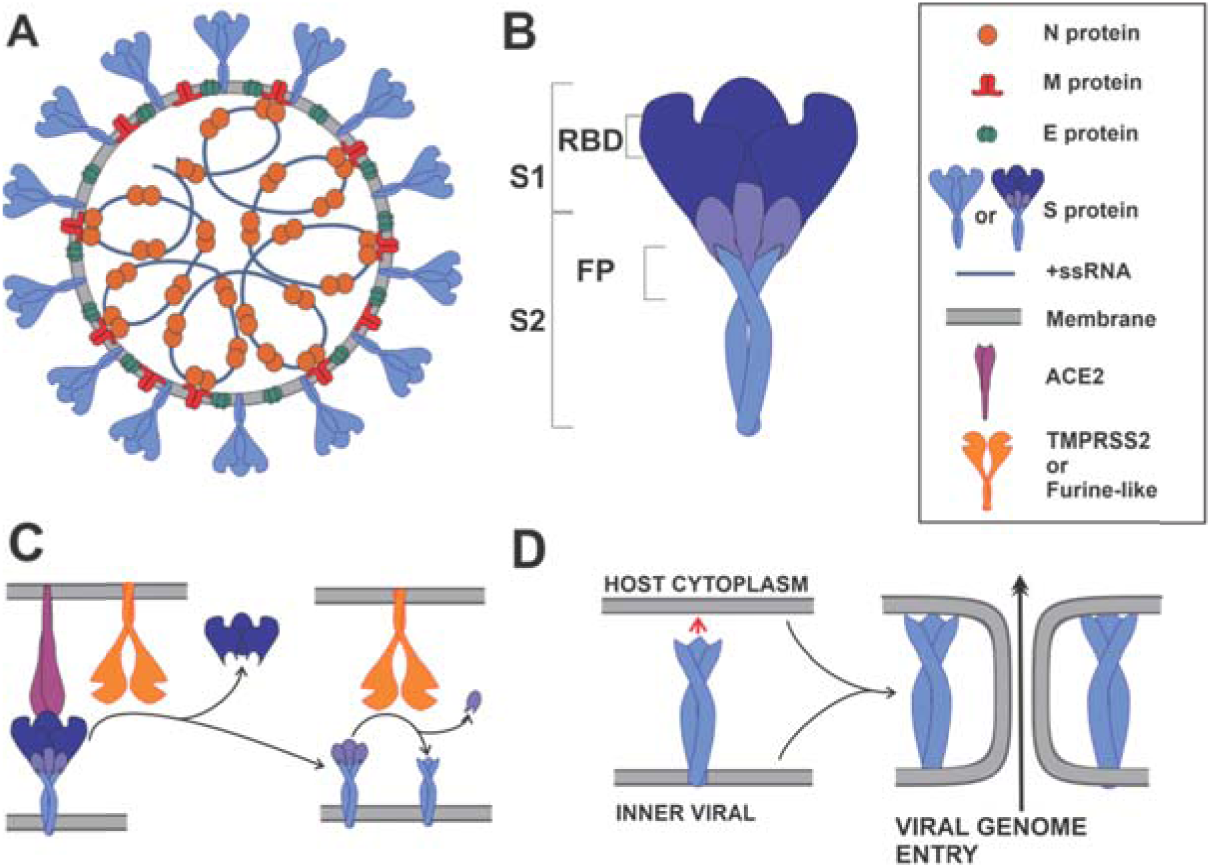
Schematic representation of a SARS-CoV-2 viral particle and key steps in virus entry. (A) The N, S, E and M proteins are represented in their oligomeric state. N protein dimers bind +ssRNA, forming the nucleocapsid. The nucleocapsid is surrounded by the viral membrane that contains S, E and M proteins. Additionally, the M protein is shown interacting with the S, E and N proteins. (B) Representative domain localization of S protein, showing the S1 and S2 fragments; S1 contains the receptor binding domain (RBD), and S2 the fusion peptide (FP). (C) Angiotensin I converting enzyme 2 (ACE2) recognition by RBD, and the subsequent S proteolytic activation by a Furine-like protease or TMPRSS2. (D) Viral and host membranes fusion induction by the exposed FP.

Once in a new host, the viral life-cycle consists basically in four stages: virus entry into a host cell, RNA translation and protein processing, RNA replication, and viral particle assembly and release [35].

During virus entry, the first stage in the viral cycle, the multidomain S protein binds to the host receptor ACE2, is proteolyzed by host proteases and activated, and thus triggers a series of events that produce the fusion of the viral membrane with the host membrane, and the subsequent release of the viral genome into the host cytoplasm [36] (Figure 1).

The second step in the virus cycle is the translation of viral structural, non-structural, and accessory proteins. Since CoVs particles do not contain any replicase protein, the translation of viral proteins is the critical step in producing all the necessary machinery for virus RNA replication and assembly. The nsp’s are translated as large (pp) pp1a and pp1b, and are cleaved by viral proteases to produce fully active proteins. CoVs posses the papain-like protease (PL^pro^), and the 3-chymotrypsin-like protease (3CL^pro^) or main protease (M^pro^). The PL^pro^ (a domain of nsp3) cleaves nsp1, nsp2 and itself, while M^pro^ (nsp5) cleaves the remaining nsp’s, including itself [37], resulting in a total of 16 nsp’s (Figure 2).

**Figure 2.**
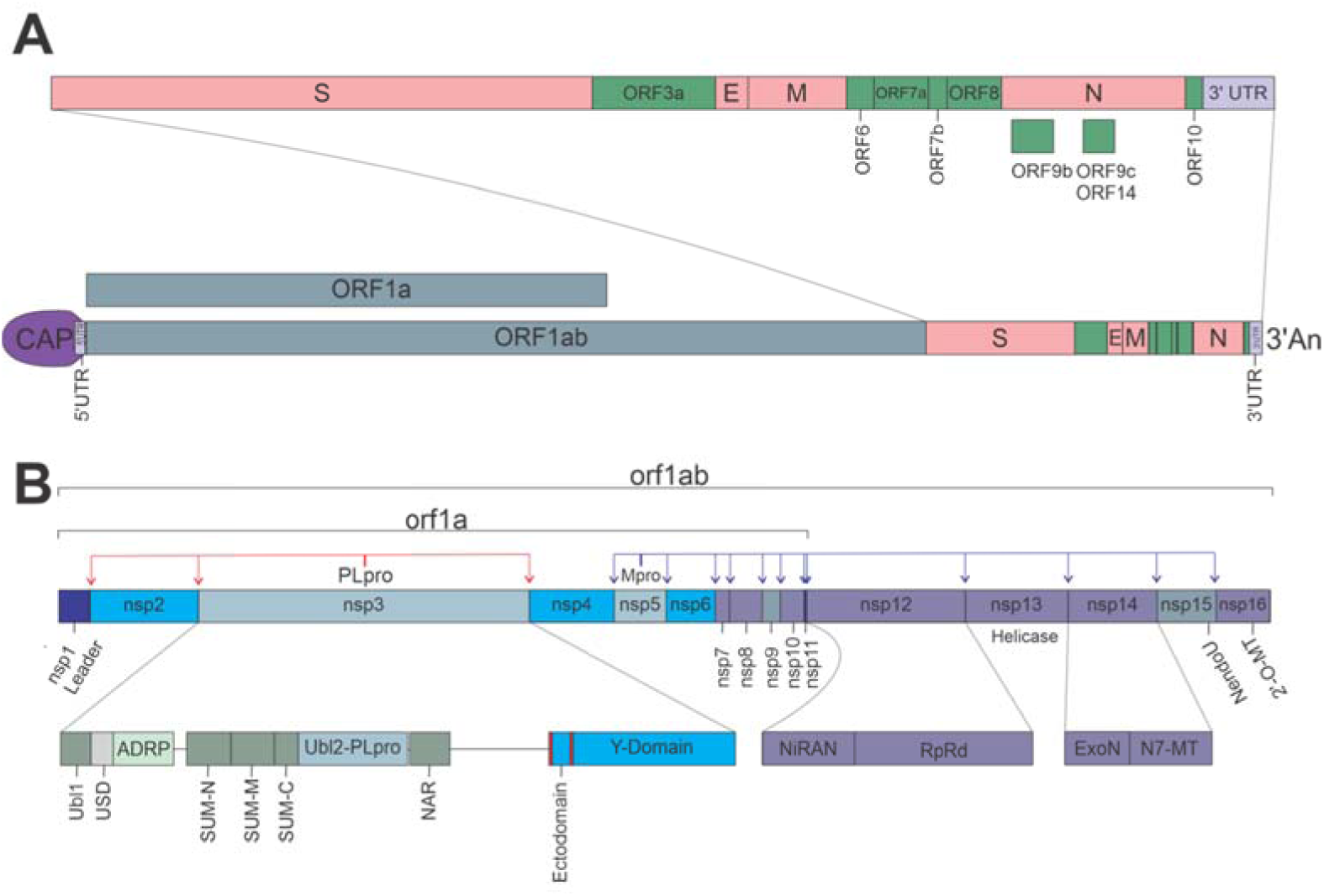
SARS-CoV2 genome organization. (A) Open reading frames (ORF) distribution in SARS-CoV-2 genome. (B) Non-structural proteins (nsp’s) distribution in orf1a and orf1ab, detailing multidomain organization of nsp3, nsp12, and nsp14; red and blue arrows indicate PL^pro^ and M^pro^ cleaving sites, respectively.

In CoVs the RNA replication takes place in double membrane vesicles (DMVs), colloquially known as viral factories [38,39]. DMVs are membrane structures derived from the Endoplasmic Reticulum (ER), formed through the action of viral membrane proteins nsp3, nsp4 and nsp6 [40]. Thanks to the ability of some nsp3 domains to bind RNA and the N protein, the viral RNA and the entire replication machin**e**ry are located in DMVs [39]. Furthermore, DMVs constitute a barrier that prevents viral RNA from being detected by the cell’s antiviral response machinery [41].

Once the DMVs are formed, the third step in the viral cycle is the replication of the viral RNA. As all +ssRNA viruses, the genome is copied to a reverse and complementary intermediate -ssRNA, which is copied back to a complete +ssRNA genome, and several shorter sub-genomic +ssRNA, that are used as additional templates to translate high amounts of structural and accessory proteins, necessary during the viral assembly [42]. CoVs require for replication the RdRp, which in this virus is the multi-protein complex nsp12-nsp8-nsp7. nsp12 has the RdRp activity, nsp8 is the putative primase, while nsp7 acts as a co-factor [43]. Since CoVs have large genomes, to achieve high efficiency and accuracy in RNA replication they possess a helicase (nsp13) that unwinds structured or double-strand RNA (dsRNA) to allow the RpRd to proceed unhindered, and a 3’-5’ exoribonuclease (ExoN, nsp14) that is capable of proofreading activity during RNA synthesis, thus lowering the rate of nucleoside misincorporation, while also enhancing resistance to nucleoside analogs [44,45] (Figure 3).

**Figure 3.**
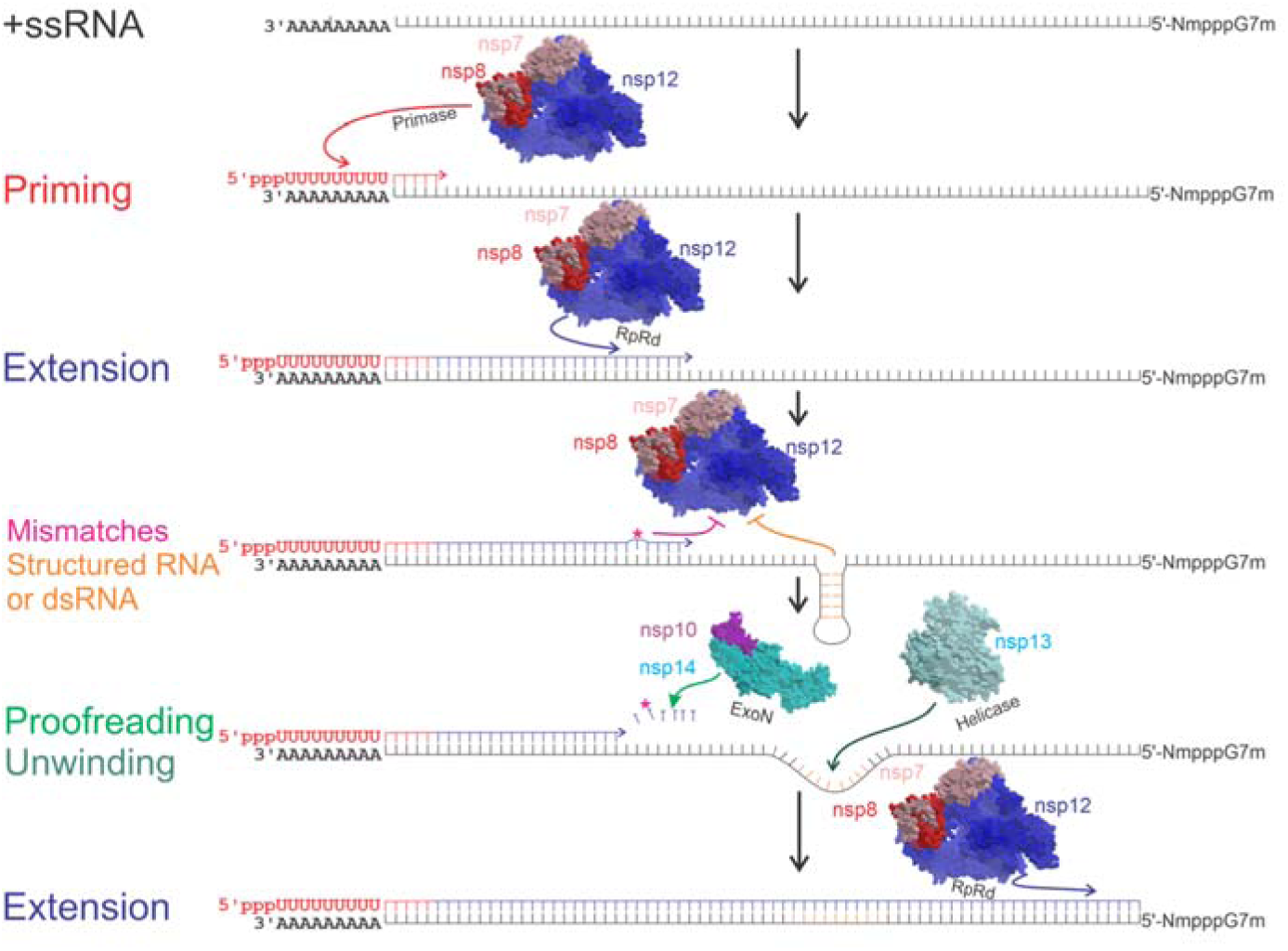
Representative steps during the synthesis of a complementary RNA strain. Priming of the complementary strain, in this case, a -ssRNA, is catalyzed by nsp8 (red letters and arrows). Primer dependent RNA extension, catalyzed by nsp7-nsp8-nsp12 complex (blue lines and arrows). Magenta asterisk (*) and arrow represent a mis-incorporated nucleotide or nucleotide analog, and nonobligate RNA chain termination, respectively. Orange lines and arrows, represent nucleotides forming dsRNA or structured RNA, and extension inhibition, respectively. Light green arrow represents nucleotide excision by nsp14 3’-5’ exonuclease (ExoN). Dark green arrow represents the unwinding activity of nsp13 helicase.

Additionally, for the viral RNA to be translated efficiently and to avoid quick degradation, CoVs have poly-adenylation [poly(A)] and capping modifications, just like host mRNAs. The poly(A) modification is apparently catalyzed by nsp8, which has TAT activity [46]. The capping consists of a sequence of reactions: 1) the terminal γ-phosphate is removed from the 5′-triphosphate end of RNA by a RNA 5’-triphosphatase (RTPase), probably nsp13; 2) an uncharacterized RNA guanylyltransferase (GTase) adds a GMP molecule to the 5’RNA, resulting in the formation of GpppN-RNA; 3) the GpppN-RNA is methylated at the N7 position of the guanosine by an N7-methyltransferase (N7-MTase, nsp14-nsp10 dimer), yielding m7GpppN (cap-0); 4) finally, a 2′-O-MTase (nsp16-nsp10 dimer), methylates the 2′-O position of the first nucleotide’s ribose of m7GpppN, yielding m7GpppNm (cap-1) [47] (Figure 4).

**Figure 4.**
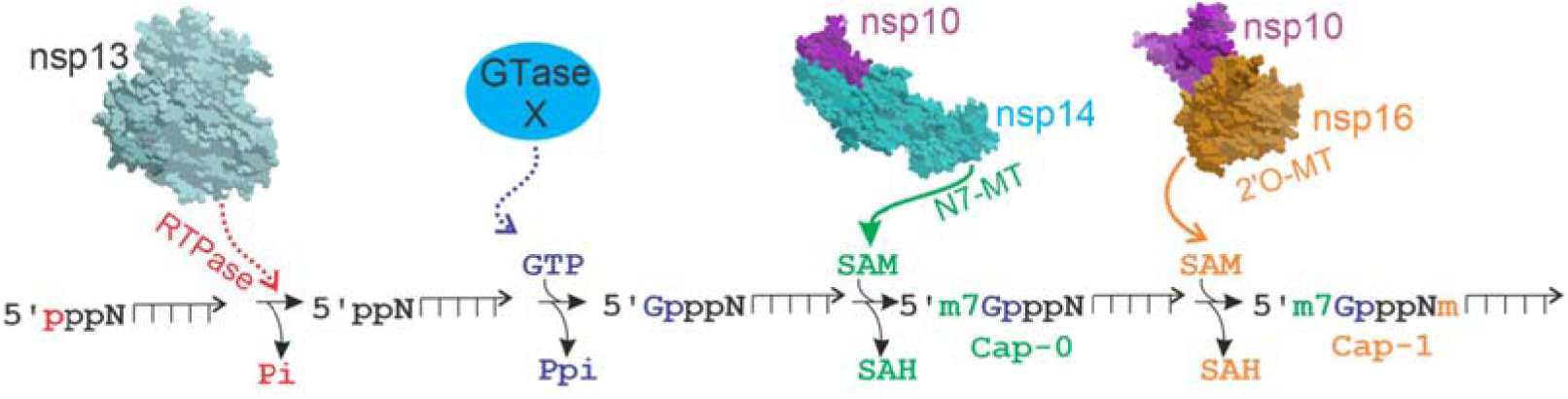
Viral RNA CAP synthesis. The capping of viral RNA has 4 enzymatic steps. i) Removal of the first phosphate of the 5′-triphosphate end of RNA by a RNA 5’-triphosphatase (RTPase), probably nsp13 (red arrow); ii) Addition of a GMP molecule to the 5’RNA by an unconfirmed guanylyltransferase (GTase) (blue); iii) Methylation of GpppN-RNA at the N7 position by nsp14 N7-methyltransferase (N7-MTase), forming the cap-0 (green arrow); iv) Methylation at the 2′-O position of the first nucleotide ribose by the 2′-O-methyltransferase (2′-O-Mtase) (nsp16), yielding the cap-1 (orange arrow). Dotted or arrow lines indicate unconfirmed enzymes. Pi: phosphate, Ppi: pyrophosphate, SAM: S-Adenosyl methionine, SAH: S-Adenosyl homocysteine, m: methyl group.

The last step in the viral cycle is the assembly of the viral particles and the release of the viral progeny. During this step, several N proteins bind and compact each copy of the genomic +ssRNA, forming the nucleocapsid. Proteins S, E, and M, accumulated in the ER-Golgi intermediate compartment (ERGIC) [48], bind the nucleocapsid, by direct M-N protein interactions [49], and then through the action of E and M proteins on the ERGIC membrane a viral particle is formed [50] in a secretory compartment [51]. Finally, the viral progeny particle or virion is released by the exocytosis pathway [2], and the viral cycle is completed.

## 3 Results

For each SARS-CoV-2 protein, we analyzed its functional role and its structural data; in cases when the latter were not available, a homology model was built, whenever possible, using the corresponding SARS-CoV structure as template. We then searched for druggable sites on the protein’s structure using FTMap, ICM Pocket Finder and CryptoSite (see Methods), and analyzed the possible functional consequences of small-molecules binding to these sites; potential allosteric sites were also identified. However, further biochemical and biological assays should be performed to confirm their actual role. We also summarized current drug repurposing efforts on the two cysteine proteases and the polymerase, but including only compounds with some kind of experimental validation.

### Non-structural protein 3 (nsp3)

Nsp3 is the largest SARS-CoV-2 protein, with 1945 aminoacids. It has at the least seven tandem macrodomains involved in different functions. These nsp3 domains are listed in Table S1, and each one is analyzed in the subsequent sections.

### The papain-like protease (PL^pro^, nsp3 domain)

The SARS-CoV-2 PL^pro^ is a domain of nsp3. It should be stressed that inhibiting viral proteases might not only be important to block the viral polyproteins processing, but since these proteases also act on host proteins -thus being partly responsible for the host cell shutdown-, it would be a way to interfere with viral immune supression. The PL^pro^ has a ubiquitin-like subdomain (Ubl2) located within its N-terminal portion, which is not necessary for PL^pro^ activity [52], and is probably involved in Nuclear Factor kappa-light-chain-enhancer of activated B cells (NFκB) signaling, affecting interferon (IFN) sensibility [53]. Through recognition of the consensus cleavage LXGG motif, PL^pro^ hydrolyzes the peptide bond on the carboxyl side of the Gly at the P1 position, thus releasing nsp1, nsp2, and nsp3 proteins. In the case of SARS-CoV PL^pro^, *in vitro* studies have also shown that this protease has two other proteolytic activities, hydrolyzing ubiquitin (Ub) and ubiquitin-like protein interferon-stimulated gene product 15 (ISG15), from cellular proteins such as Interferon regulatory factor 3 (IRF3), thus suppressing the host innate immune response [54-57].

The crystal structures of SARS-CoV-2 PL^pro^ in complex with inhibitors VIR250 and VIR251 covalently attached to the catalytic C111 were solved at 2.8 Å (VIR250, PDB 6WUU) and 1.65 Å (VIR251, PDB 6WX4), respectively (Figures S1 and 5a). These inhibitors displayed similar activities towards SARS-CoV and SARS-CoV-2 PL^pro^s, but a weaker activity towards the MERS-CoV PL^pro^. Two wild type apo structures (PDBs 6W9C and 6WZU, at 2.7 Å and 1.8 Å, respectively), and the C111S apo structure (PDB 6WRH, 1.6 Å) have overall main chain RMSD values of ∼0.6 Å compared to the lowest resolution PL^pro^-VIR251 structure, displaying a partial collapse of the catalytic binding site. SARS-CoV-2 PL^pro^ was crystallized also in complex with Ub (PDB 6XAA) at 2.7 Å, and with the C-terminal part of ISG15 (PDB 6XA9) at 2.9 Å (Figure S1). Both structures are very similar to 6XW4, with main chain RMSD values of ∼0.5 Å (all available PL^pro^ structures are listed in Table S1).

In agreement with another work [58], it was shown that while SARS-CoV-2 PL^pro^ retained de-ISGylating activity similar to its SARS-CoV counterpart, its hydrolyzing ability of diUb^K48^ was decreased [59]; this might not be surprising, since specificity for Ub and ISG15 substrates might depend on only one mutation [60-62], and while the Ub S1 site is conserved between the PL^pro^s of SARS-CoV and SARS-CoV-2 (83% similarity, 17% identity), the Ub S2 site has only 67% similarity and 13% identity. Interestingly, SARS-CoV-2 PL^pro^ displays preference for ISG15s from certain species including humans [58]. It has yet to be determined whether this functional difference has any implication in the ability of SARS-CoV-2 to evade the human innate immune system.

The catalytic site (Figure 5) would clearly the first choice for drug discovery (see below successful cases targeting this site). Using FTMap, a borderline druggable binding site delimited by residues R166, L185, L199, V202, E203, M206-M208, and K232, was identified; a small-molecule binding to this site might interfere with Ub binding, but not with ISG15 (Figure 5b). A small hot spot could also be found limited by amino acids P59, R65, and T74-F79. This site is close to the Ub S2 binding site, and from the structural model of the SARS-CoV-2 PL^pro^-diUb^K48^ [generated using the corresponding SARS-CoV structure (PDB 5E6J)], a small-molecule binding to it might interfere with Ub binding. Two potential binding sites were identified in PL^pro^, delimited by residues: i) S213-E215, K218, Y252, L254, K255, T258-T260, V304, Y306, and E308; ii) L121-I124, L126-F128, L133, Q134, Y137, Y138, and R141 (Figure S1). These sites are far from the catalytic, Ub1-, and Ub2-binding sites, so even in the case a small-molecule could bind to them, further evidence is needed to determine whether those amino acids have a functional role, and to assess if those sites could display allosteric modulation potential.

**Figure 5.**
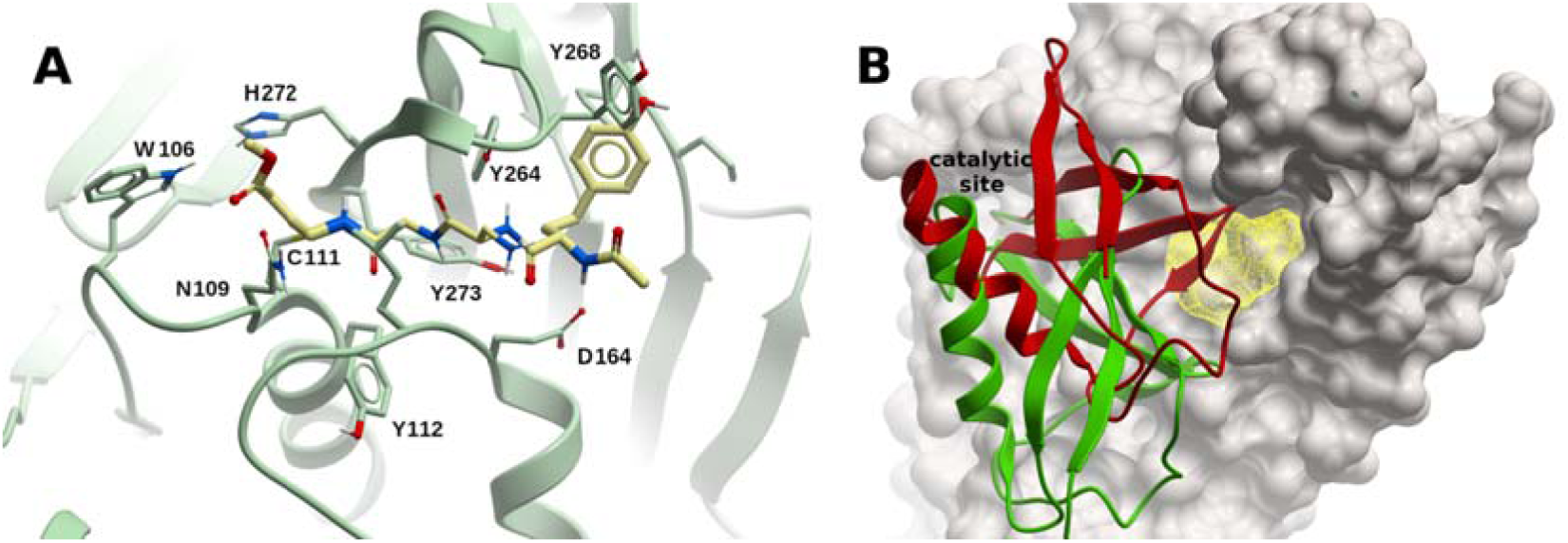
Structure and binding sites of SARS-CoV-2 PL^pro^. (A) Complex of PL^pro^ with inhibitor VIR251 covalently bound to C111 within the catalytic binding site (PDB 6XW4). (B) Potential binding site (yellow mesh) on SARS-CoV-2 PL^pro^ (ivory surface). Ubiquitin (red) and ISG15 (green) are displayed in ribbon representation. The N-terminus of these two proteins are inserted within the catalytic site. A small-molecule binding to the predicted site would interfere with ubiquitin binding, but not with ISG15.

Considering that the residues lining the catalytic sites of SARS-CoV and SARS-CoV-2 PL^pro^s are identical, Freitas et al. evaluated naphtalene-based SARS-CoV non-covalent inhibitors on SARS-CoV-2, and found that two compounds, GRL-0617 and **6**, exhibited IC_50_ values of 2.4 μM and 5.0 μM, respectively (their corresponding values in SARS-CoV PL^pro^ were 600 nM and 2.6 μM), displaying in antiviral activity assessment EC_50_ values of 27.6 and 21.0 μM, respectively, with no cytotoxicity in cell cultures [58]. The characterization of the non-covalent inhibition between these ligands and SARS-CoV-2 PL^pro^ was performed using a homology model based on SARS-CoV PL^pro^ bound to GRL-0617 (PDB 3E9S, [63]). It should be stressed that a similar approach using MERS-CoV inhibitors [64] would not be so straightforward, considering the lower conservation of the catalytic binding site at the sequence level. It was also shown that the synthetic organoselenium drug molecule ebselen, which displays anti-inflammatory, anti-oxidant and cytoprotective activity in mammalian cells, covalently inhibits the enzymatic activity of SARS-CoV-2 PL^pro^, with an IC_50_ ∼2 μM, exhibiting a weaker activity against its SARS-CoV counterpart [65]. In a follow up contribution, ebselen-derivatives were identified, displaying lower inhibition constants, in the range of 250 nM [66]. Klemm et al. showed that benzodioxolane analogs **3j, 3k**, and **5c**, which inhibit SARS-CoV PL^pro^ in the sub-micromolar range [67] displayed similar inhibitory activity against SARS-CoV-2 PL^pro^ [68]. Two other compounds have shown to inhibit SARS-CoV-2 PL^pro^ *in vitro*, inhibiting viral production in cell culture, namely, the approved chemotherapy agent 6-Thioguanine [69], and the anti-dengue protease inhibitor mycophenolic acid [70]. The dual viral polypeptide cleavage and immune suppression roles of PL^pro^ make it an attractive target for antiviral development.

### Main protease (M^pro^, nsp5)

The SARS-CoV-2 M^pro^ is a cysteine protease involved in most cleavage events within the precursor polyproteins, beginning with the autolytic cleavage of itself from pp1a and pp1ab [71] (Figure 2). The vital functional role of M^pro^ in the viral life cycle, coupled with the absence of close human homologs, makes M^pro^ an attractive antiviral target [72-76].

The active form of M^pro^ is a homodimer containing two protomers composed of three domains each [77]: domain I (residues F8–Y101), domain II (residues K102–P184), and domain III (residues T201–V303); domains II and III are connected by a long loop region (residues F185–I200). The M^pro^ has a Cys-His catalytic dyad (C145-H41), and the substrate binding site is located in the cleft between domains I and II (Figure S2). The superposition of 12 crystal structures of M^pro^ from different species [77] showed that the helical domain III and surface loops display the largest conformational variations, while the substrate-binding pocket is highly conserved among M^pro^ in CoVs. This confirmed an earlier hypothesis that the substrate-recognition site was highly conserved across CoVs, what could serve to design pan-CoV inhibitors [76].

Crystal structures of SARS-CoV-2 M^pro^ were determined in complex with the Michael acceptor covalent inhibitor N3 (Figure 6a) at 2.1 Å resolution (PDB 6LU7), and 1.7 Å (PDB 7BQY), respectively (an up-to-date detailed list of M^pro^ solved structures is shown in Table S2). N3 inhibits the M^pro^ from multiple CoVs, and has been co-crystalized with M^pro^ in SARS-CoV (PDB 2AMQ), Infectious Bronchitis Virus (IBV) (PDB 2Q6F), Human coronavirus HKU1 (HCoV-HKU1) (PDB 3D23), Feline Infectious Peritonitis Virus (FIPV) (PDB 5EU8), Human coronavirus NL63 (HCoV-NL63) (PDB 5GWY), Porcine Epidemic Diarrhea Virus (PEDV) (PDB 5GWZ), and Mouse Hepatitis Virus (MHV) (PDB 6JIJ). In the SARS-CoV-2 M^pro^-N3 complexes, the Sγ atom of C145 forms a covalent bond with the Cβ atom of the vinyl group of N3 (see Table S2 for available structures solved in the apo form, and with covalent and non-covalent inhibitors; more than hundred crystal structures with bound fragment-like molecules are not included, cf. Ref. [78]).

**Figure 6.**
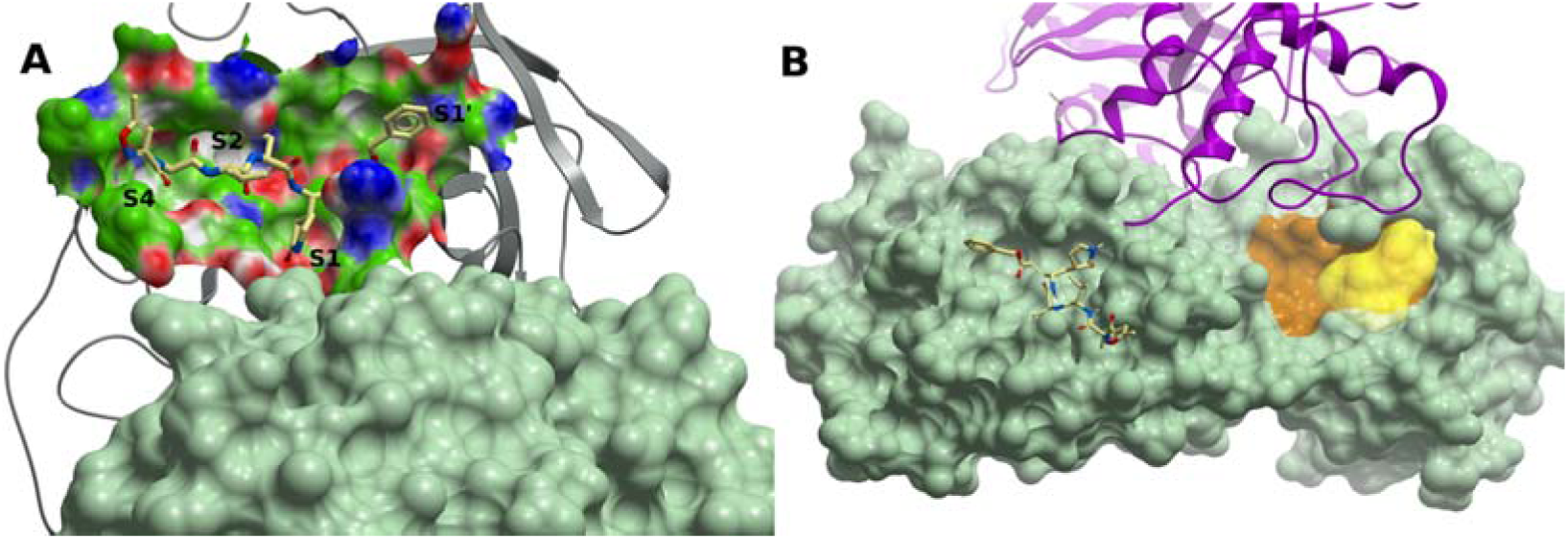
SARS-CoV-2 main protease M^pro^ binding sites. (A) M^pro^ (grey ribbon) in complex with peptide inhibitor N3 (PDB 6LU7). The protein subsites S1, S2, S4 and S1’ are labeled. The molecular surface of the other protomer is shown in light green. (B) A cryptic site on M^pro^ (brown) with a CS site nearby lined up by residues T199, Y237, Y239, L271, L272, G275, M276, and A285-L287 (yellow surface). M^pro^ is represented by a green molecular surface. N3 is also displayed within the catalytic site (light yellow carbons). The other protomer is represented in magenta ribbon. A small-molecule binding to this potential site might interfere with homodimerization.

The primary choice for drug discovery would be the catalytic site. Zhang et al. crystallized SARS-CoV-2 M^pro^ in its apo form (PDB 6Y2E) to guide the design of a novel α-ketoamide inhibitor displaying an IC_50_ = 0.67 μM; crystal structures of SARS-CoV-2 in complex with this inhibitor were determined in the monoclinic and orthorombic form (PDBs 6Y2F and 6Y2G, respectively) [79]. The structure of M^pro^ in complex with antineoplastic drug carmofur, which inhibits viral replication in cells with EC_50_ = 24.3 μM, was determined at 1.6 Å resolution [80]; the carbonyl group of carmofur attaches covalently to catalytic C145 and the fatty acid tail occupies the S2 subsite. Anti-inflammatory ebselen has also shown a promising inhibitory effect in M^pro^ and reduction of the virus titer in cell culture [77]. Starting with the substrate-binding site of the SARS-CoV M^pro^, Dai et al. designed and synthesized two novel covalent inhibitors of the SARS-CoV-2 M^pro^, displaying anti-SARS-CoV-2-infection activity in cell culture with EC_50_ values of 0.53 μM and 0.72 μM, respectively, with no significant cytotoxicity [81]. Some known protease inhibitors have been identified *in silico* and tested *in vitro* on M^pro^, such as the HIV protease inhibitors ritonavir, nelfinavir, saquinavir, and atazanavir [82,83]; the hepatitis C virus (HCV) protease inhibitor boceprevir, the broad-spectrum protease inhibitor GC-376, and calpain inhibitors II and XII were also shown to inhibit viral replication by targeting the M^pro^ [84].

A cryptic site with a CS within 5 Å defined by residues T199, Y237, Y239, L271, L272, G275, M276, and A285-L287 was identified; this site is not too distant from the partner protomer, thus it should be explored whether a molecule binding to it might preclude dimer formation (Figure 6b). A second borderline druggable site delimited by residues Q107-Q110, V202, N203, H246, I249, and T292-F294 was also identified. This site lies on the opposite side of the dimerization interface, and the functional consequences of binding to it have yet to be determined.

### The RNA polymerase complex (nsp12-nsp7-nsp8)

In +ssRNA viruses, the synthesis of RNA is catalyzed by the RpRd, in a primer-dependent manner [85]. In SARS-CoV-2 and other CoVs, the RpRd complex consists of a catalytic subunit (nsp12), and co-factors nsp7 and nsp8 [86,87].

The SARS-CoV-2 RdRp experimental structures contain two nsp8 molecules: one forming a heterodimer with nsp7 (nsp8-2), and a second one bound at a different site (nsp8-1), in a similar fashion as in SARS-CoV [87] (cf. Table S3 for a list of the available RdRp complex experimental structures). Interaction with nsp7 and nsp8 provides stability to nsp12 [43], consistent with the observation that nsp12 in isolation displays little activity, while the presence of nsp7 and nsp8 enhance template binding and processivity [87-89], as do other DNA/RNA-binding proteins associated to a polymerase, such as the Proliferating Cell Nuclear Antigen (PCNA). Recently, the crystal structure of RdRp in complex with the helicase (nsp13) has also become available (PDB 6XEZ) [90].

The overall structure of the SARS-CoV-2 RdRp complex is very similar to that of SARS-CoV, with a Cα RMSD of ∼0.8 Å, consistent with the high degree of sequence similarity (nsp7, 99%; nsp8, 97%; nsp12, 97%). Although the amino acid substitutions are not located in the catalytic site, the SARS-CoV-2 RdRp complex displays a 35% lower efficiency for RNA synthesis than its SARS-CoV counterpart; in fact, this lower efficiency is due to changes restricted to nsp8 and nsp12, not nsp7 [43].

The nsp12 contains a right-hand RdRp domain (residues L366-F932) −a conserved architecture of viral RdRps− and a N-terminal nido-virus RdRp-associated nucleotidyl-transferase (NiRAN) domain (residues D51-R249); these two domains are linked by an interface domain (residues A250 to R365). The polymerase domain is formed by three subdomains: a fingers subdomain (residues L366-A581, and K621-G679), a palm subdomain (residues T582-P620 and T680-Q815), and a thumb subdomain (residues H816-E932) (Figure 7a). An N-terminal β-hairpin (residues D29-K50) establishes close contacts with the NiRAN domain and the palm subdomain, and contributes to stabilize the structure. As in the corresponding SARS-CoV nsp12 [87], the integrity of the overall structure is maintained by two zinc ions that are present within metal-binding sites composed of H295-C301-C306-C310 and C487-H642-C645-C646.

**Figure 7.**
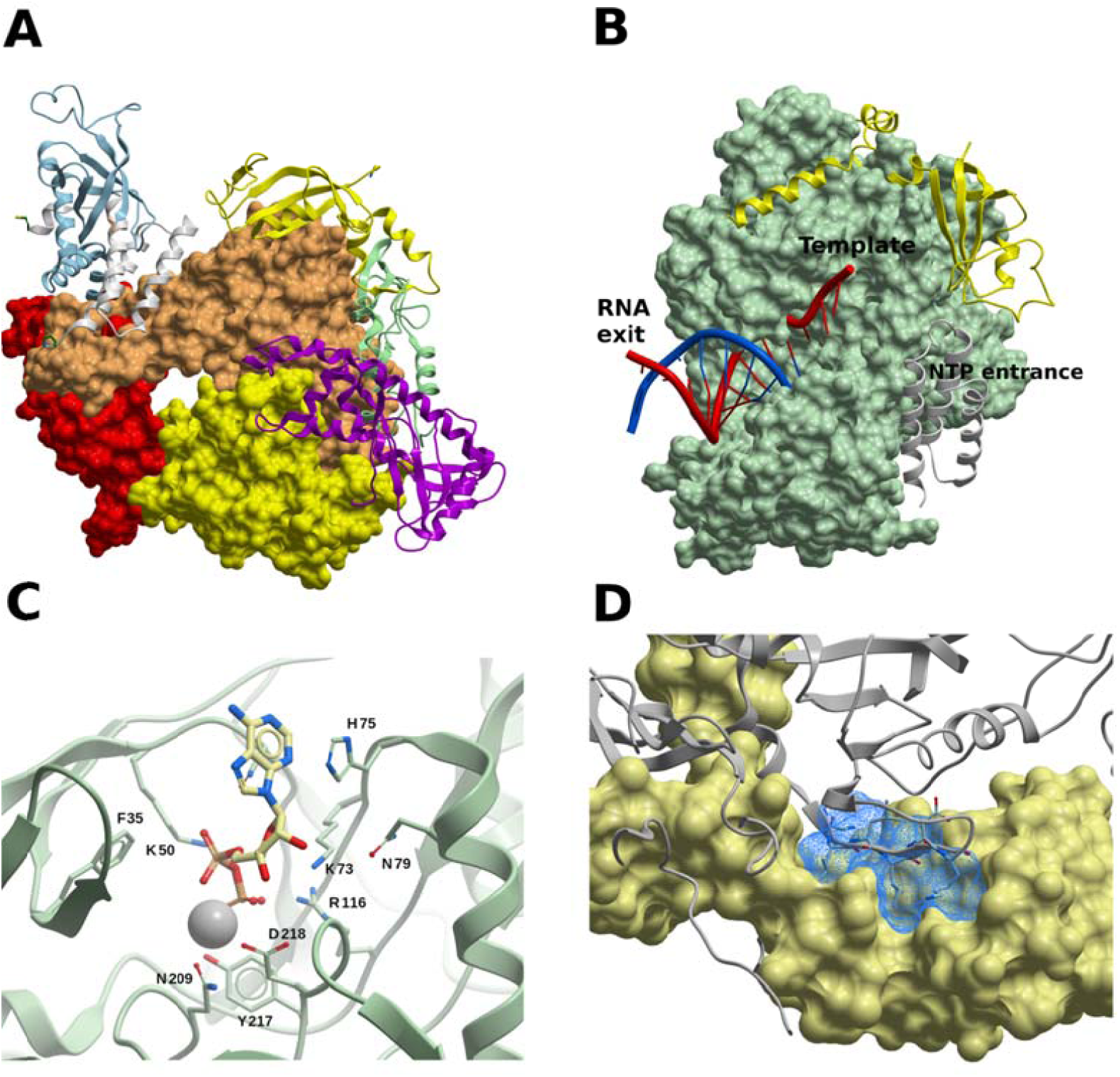
The SARS-CoV-2 RdRp complex (nsp12-nsp8-nsp7). (A) Structure of the nsp12-nsp7-nsp8 complex. The channel in the middle corresponds to the catalytic active site. Color code: nsp7, white ribbon; nsp8-2, light blue ribbon; nsp8-1, yellow ribbon. The nsp12 domains are colored as follows: palm, yellow; fingers, tans; palm, red; interface, pale green ribbon; NiRAN, magenta ribbon. (B) RdRp complex bound to RNA. Nsp12 is displayed as a green surface. Color code: Primer RNA, blue; template RNA, red; nsp7, yellow ribbon; nsp8-1, grey ribbon. (C) Molecule of ADP-Mg^2+^ within the NiRAN domain of nsp12. Interacting residues are shown, in what may constitute a druggable binding site. (D) Target site in nsp8 (light yellow surface). The predicted binding site is represented using a blue mesh representation, and nsp12 is shown as gray ribbon. Nsp12 residues N386-K391 are displayed (though not labeled) to highlight that a small molecule binding to these potential sites might interfere with PPIs.

The RdRp-RNA structures show that RNA mainly interacts with nsp12 through its phosphate-ribose backbone, especially involving the 2’-OH groups. No base pair-protein contacts are observed, pointing at sequence-independent binding of RNA and enzymatic activity. Except for three specific and localized structural changes [89], the overall structures of the apo and RNA-bound complexes are very similar, displaying a main chain RMSD value of ∼0.5 Å; the absence of relevant conformational rearrangements from the apo state implies that the enzyme can start to function as replicase upon RNA binding, which also correlates with its high processivity. The catalytic active site of the SARS-CoV-2 RdRp is composed by the seven conserved polymerase motifs A-G in the palm domain, as in other RNA polymerases: Motif A, P612-M626; B, G678-T710; C, F753-N767; D, A771-E796; E, H810-V820; F, L544-R555; G, K500-L514. These motifs and the RNA binding residues are highly conserved across viral polymerases [91]. It should be noted that in RdRp structures with longer RNA products (PDBs 6YYT, 26 bases, and 6XEZ, 34 bases), the long α-helical extensions of both nsp8 molecules form positively charged ‘sliding poles’ along the exiting RNA, in what could be a supporting scaffold for processing the large CoV genome. Since these nsp8 extensions are mobile in free RdRp [86,87], they may adopt an ordered conformation upon dsRNA exit from nsp12.

Remdesivir, a drug designed for Ebola virus [92], has received a great deal of attention as a RdRp inhibitor and potential treatment for COVID-19 [93]. Remdesivir is a prodrug that is converted to the active drug in the triphosphate form [remdesivir triphosphate (RTP)] by host enzymes. Two cryo– electron microscopy structures of the SARS-CoV-2 RdRp complex have been solved, one in the apo form, and the other in complex with a template-primer RNA and with remdesivir covalently bound to the primer strand [89] (PDBs 7BV1 and 7BV2, respectively) (Figure 7b). At high RTP concentration, the mono-phosphate form of remdesivir (RMP) is added (at position i) into the primer-strand [89], thus causing termination of RNA synthesis at position i+3, in SARS-CoV-2, SARS-CoV, and MERS-CoV [94]. Favipiravir (also called avifavir), used in Hepatitis C Virus (HCV) treatment, are also promising drugs [93]. Nevertheless, other antivirals, such as sofosbuvir, alovudine, tenofovir alafenamide, AZT, abacavir, lamivudine, emtricitabine, carbovir, ganciclovir, stavudine, and entecavir, are incorporated by SARS-CoV-2 RdRp and block replication *in vitro* [95,96]. Another type of broad spectrum ribonucleoside analog, β-D-N4-hydroxycytidine (EIDD-1931), has been shown to inhibit SARS-CoV-2, SARS-CoV and MERS-CoV in cell culture, by increasing the mutation transition rate, probably exceeding the proofreading ability granted by ExoN [97].

In the RdRp-nsp13 complex (PDB 6XEZ), the NiRAN domain is occupied by an ADP-Mg^2+^ molecule (Figure 7c). While the target of the NiRAN nucleotidyltransferase activity is unknown, this activity itself is essential for viral propagation [98]. Thus, this binding site might constitute an interesting novel druggable site in nsp12, and it should be further assessed how a small-molecule binding to it would interfere with the viral cycle.

The design of non-nucleoside inhibitors calls for the analysis of alternative disruptive sites within the RdRp complex, considering that the assembly of the nsp12-nsp7-nsp8 complex is needed for RNA synthesis. We used FTMap to identify druggable sites on PPI interfaces in nsp12, nsp7, and nsp8. Several hot spots (consensus sites) were identified on nsp8, three of which formed a potential druggable site, outlined by residues P121, A125, K127-P133, T137, T141, F147, and W154 (Figure 7d). This site lies within the nsp12-nsp8 interface, and a molecule binding to this site could interfere with the interaction of the β-strand N386-K391 of nsp12 with nsp8 (Figure 7d). A borderline druggable target site was identified on nsp12, lined up by residues L270, L271, P323, T324, F326, P328-V330, R349, F396, V398, M666, and V675 (Figure S3), where a molecule binding to it might interfere with nsp8 binding by clashing with its segment V115-I119. On nsp7, a borderline druggable site was identified defined by residues K2, D5, V6, T9, S10, V12, F49, and M52-S54, within the PPI interface with nsp8-2.

In addition to its functionality within the RdRp complex, the nsp7-nsp8 hexadecamer has *de novo* initiation of RNA synthesis capability also known as primase activity, while nsp8 has also displayed TATase activity. Analysis of the SARS-CoV nsp7-nsp8 hexadecameric structure (PDB 2AHM [99]) shows several sites which could be targeted to interfere with its primase activity, making impossible for nsp12 to extend the complementary strand due to the lack of primer. Moreover, there is a possibility that small-molecules could interfere with the conformational dynamics needed to interact within the RdRp complex. However, further structural and computational studies (such as molecular dynamics, MD) would be needed to confirm this hypotheses.

### Helicase (nsp13)

The SARS-CoV-2 nsp13 possesses helicase activity, thus playing a key role in catalyzing the unwinding of dsRNAs or structured RNAs in the 5’ to 3’ direction into single strands. It has been demonstrated that in SARS-CoV this happens in an NTP-dependent manner [45,100]. Additionally, it has been shown that the helicase has a RTPase activity (which may be the first step for the formation of the 5’ cap structure of viral RNAs), and that the nsp13-associated NTPase and RNA 5’-triphosphatase activities share a common active site, both for SARS-CoV [101] and the human CoV 229E (HCoV-229E) [102].

The crystal structure of the SARS-CoV-2 nsp13 has been solved at 1.9 Å (PDB 6ZSL). A structure solved by cryo-EM of nsp13 in complex with RdRp (nsp12-nsp8-nsp7) is also available (PDB 6XEZ [90]). Nsp13 has the form of a triangular pyramid with five domains. At the top, the N-terminal zinc binding domain with three zinc fingers (A1-S100) is connected with the stalk domain (D101-G150); then, at the base of the pyramid, three domains (1B, I151-E261; 1A, F262-R442; 2A, R443-N596) form the triangular base (Figure S4). Compared to its SARS-CoV and MERS-CoV counterparts, the SARS-CoV-2 nsp13 shares a 99.8% (100%), and 71% (82%) sequence identity (similarity), respectively. The nsp13 structures are also very similar, with backbone RMSD values of ∼1.9 Å between SARS-CoV-2 nsp13 and the corresponding SARS-CoV (PDB 6JYT) [45] and MERS-CoV (PDB 5WWP) [103].

In the SARS-CoV nsp13, six residues were identified as being involved in NTP hydrolysis (K288, S289, D374, E375, Q404 and R567), and mutations of any of these residues to alanine resulted in high unwinding deficiency and decreased ATPase activity [45]. Moreover, it was also shown in the same study that, for all six mutants, changes in helicase activity are consistent with changes in their ATPase activity, thus demonstrating that nsp13 performs its helicase activity in an NTPase-dependent manner. As stated above, this site would also correspond to the RTPase activity. The SARS-CoV-2 nsp13 structure PDB 6XEZ features an ADP-Mg^2+^ molecule in the vicinity of those residues (Figure 8a), suggesting that targeting this site may interfere with the NTPase or RTPase functions.

**Figure 8.**
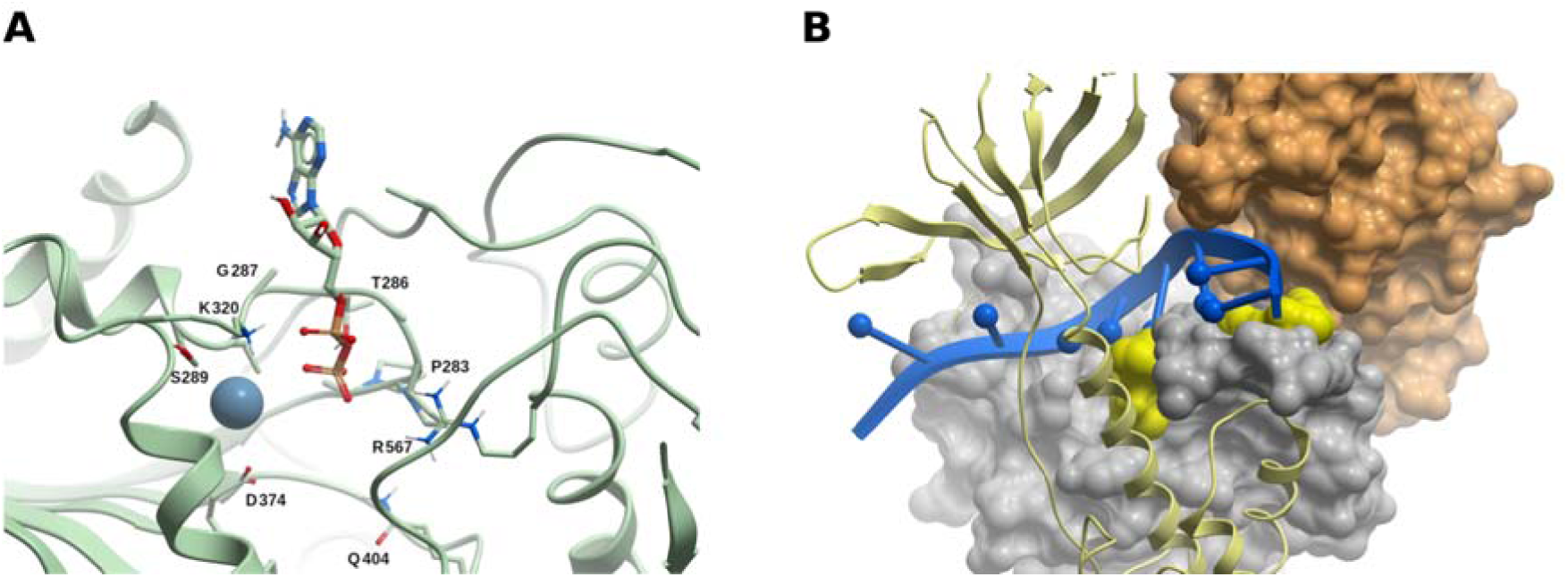
The SARS-CoV-2 helicase/NTPase/RTPase (nsp13). (A) The ADP-Mg^2+^ bound within nsp13. This site has been identified as being involved in NTP hydrolysis in SARS-CoV, and cou**l**d constitute a druggable site. (B) Two potential borderline druggable binding sites identified in nsp13. The structure of RNA (blue) was modeled based on the yeast Upf1-RNA complex structure (PDB 2XZL). Nsp13 is represented as a green ribbon, but nsp13 domains 1A and 2A are displayed as grey and tan molecular surfaces, respectively. The potential sites are shown in yellow. Based on our model, molecules binding to these sites might interfere with RNA binding.

The zinc binding and stalk domains are important for the helicase activity in SARS-CoV [45]. Mutations to alanine of the zinc binding domain residue N102 (which interacts with the stalk domain) and of the stalk residue K131 (which interacts with the 1A domain) resulted in decreased helicase activity. This appears to be a top-to-bottom signaling system, and it is difficult to figure out a way to interfere in this process with small molecules, with the information available up to today. The zinc binding domain might be targeted with metal chelators such as bismuth complexes.

Based on an interaction model of dsRNA with SARS-CoV nsp13, it was hypothesized that residues 176–186 (1B domain), 209–214 (1B domain), 330–350 (1A domain) and 516–541 (2A domain) constitute a probable nucleic acid binding region [45]. While there is no CoV nsp13 structure with a nucleic acid substrate bound, we performed a crude model of SARS-CoV-2 nsp13 in complex with RNA based on a Yeast-Upf1-RNA structure (PDB 2XZL), in a similar fashion as in an earlier work [45], and in agreement with a recent model [90] (Figure 8b). Using FTMap, two potential borderline druggable sites were identified delimited by amino acids P406-409, L412, T413, G415, L417, F422, N557, and R560, and K139, E142, C309, M378-D383, P408, and T410, which could interfere with RNA binding, according to our artificial model (Figure 8b).

### Exoribonuclease/guanine-N7 methyl transferase (nsp14-nsp10 complex)

While the RdRp complex (nsp12-nsp8-nsp7) allows the synthesis of large viral RNAs due to an increased processivity conferred by the heteromeric complex, to achieve high accuracy, CoVs exhibit 3’-5’ exoribonuclease (ExoN) activity for proofreading of the viral genome synthesis, located in the N-terminal domain of nsp14. The ExoN excises mutagenic nucleotides misincorporated by the RdRp, thus conferring potential drug resistance to nucleoside analogs inhibitors. It has been reported that the ExoN activity protects SARS-CoV from the effect of base analog 5-fluorouracil [104], and that guanosine analog ribavirin (Rbv) 5’-monophosphate is incorporated at the 3’-end of RNA by the SARS-CoV RdRp, but excised from RNA by the nsp14-nsp10 ExoN (to a lesser degree by the nsp14-ExoN alone), which could account for the poor effect of Rbv in treating CoV-infected patients [105,106]. Since it was shown that SARS-CoV displays a significantly lower nucleotide insertion fidelity *in vitro* than that of the Dengue RdRp [107], it could be concluded that the low mutation rate of SARS-CoV is related to the ExoN activity. This clearly shows that targeting the ExoN function could be an excellent strategy for the development of pan-CoV therapeutics, complementing existing RdRp inhibitors, like Remdesivir. In fact, MHV lacking the ExoN activity was shown to be more susceptible to Remdesivir [108].

The C-terminal domain of nsp14 functions as a guanine-N7 methyl transferase (N7-MTase), with S-adenosyl-L-methionine (SAM) being demethylated to produce S-adenosyl-L-homocysteine (SAH), transferring the methyl group at the N7 position of guanine in 5’GpppN in viral RNAs, thus forming m7GpppN (cap-0). This capping is followed by methylation of the ribose of the first nucleotide at the 2’-O position by the 2’-O-MTase (nsp16), forming m7GpppNm (cap-1) (Figure 4). The capping structure is a protective and pro-transductional modification of the viral RNA. Firstly, the capping protects the viral RNAs from being susceptible to the 5’-3’ exoribonucleases (Xrn’s), a process called 5’ RNA decay [109]; secondly, translation initiation factors, such as the eukaryotic translation initiation factor 4E (eIF4E), binds to the cap, being this binding the limitating reaction to the load of ribosomes to the viral RNA, and also for RNA circularization; these two activities are necessary for higher translation rates due the ribosome recycling [110]. That is why blocking MTase activity may increase viral RNA decay and suppress viral RNA translation.

While there is no experimental structure available of SARS-Cov-2 nsp14, crystal structures of the SARS-CoV nsp10-nsp14 dimer are available [107,111], with nsp14 in complex with SAM at 3.2 Å (PDB 5C8T), with SAH and guanosine-P3-adenosine-5′,5′-triphosphate (GpppA) at 3.3 Å (PDB 5C8S), and in its unbound form (PDBs 5C8U and 5NFY, both at 3.4 Å). The sequence identity between SARS-Cov-2 and SARS-CoV nsp14 is 95% (similarity 98%), with no gaps in the alignment and full conservation in all functionally relevant sites. SARS-CoV nsp10 and its SARS-CoV-2 counterpart share 98% sequence identity (similarity 100%). We thus built a homology model of the SARS-COV-2 nsp10-nsp14 dimer using the corresponding SARS structure 5C8T as template (see Methods); the missing S454-D464 segment within the template was included in the model, and optimized. Since there are no gaps, the numbering scheme of template and model coincides.

In nsp14, amino acids A1-C285 fold into the ExoN domain, and the N7-MTase function lies within amino acids D301-Q527; both domains are connected by a loop (amino acids F286-G300), whose abolition was shown to suppress the N7-MTase function in SARS-CoV [112] (Figure 9a). The architecture of the catalytic core and active sites of the ExoN domain are similar to those of the DEDD superfamily exonucleases, though exhibiting some differences [107,111]. The catalytic residues D90, E92, E191, D273 and H268 (DEEDh motif) display similar structural arrangements to other proofreading homologs, such as the DNA polymerase I (1KLN) and the ε subunit of DNA polymerase III of *E. coli* (1J53), but with only a single Mg^2+^ ion at its active center. Mutating any of these residues to alanine either impaired the ExoN activity or severely reduced the ability to degrade RNA in SARS-CoV [111]. The ExoN domain also contains two zinc fingers, the first one comprising residues C207, C210, C226 and H229, and the second one in proximity to the catalytic site, comprising residues H257, C261, H264 and C279. In SARS-CoV, none of the mutants of zinc finger 1 could be expressed as soluble proteins, thus revealing its importance in protein stability, and mutants of zinc finger 2 had their enzymatic activity abolished [111]. The ExoN domain interacts with nsp10 (Figure 9a), exhibiting an interaction surface of ∼7,800 Å^2^, more than twice the nsp10-nsp16 complex. It has been reported that SARS-CoV nsp10 is necessary for the correct positioning of the residues of the ExoN catalytic site, which partially collapses in the absence of nsp10, and thus explains the reduced ExoN activity of nsp14 alone [44].

**Figure 9.**
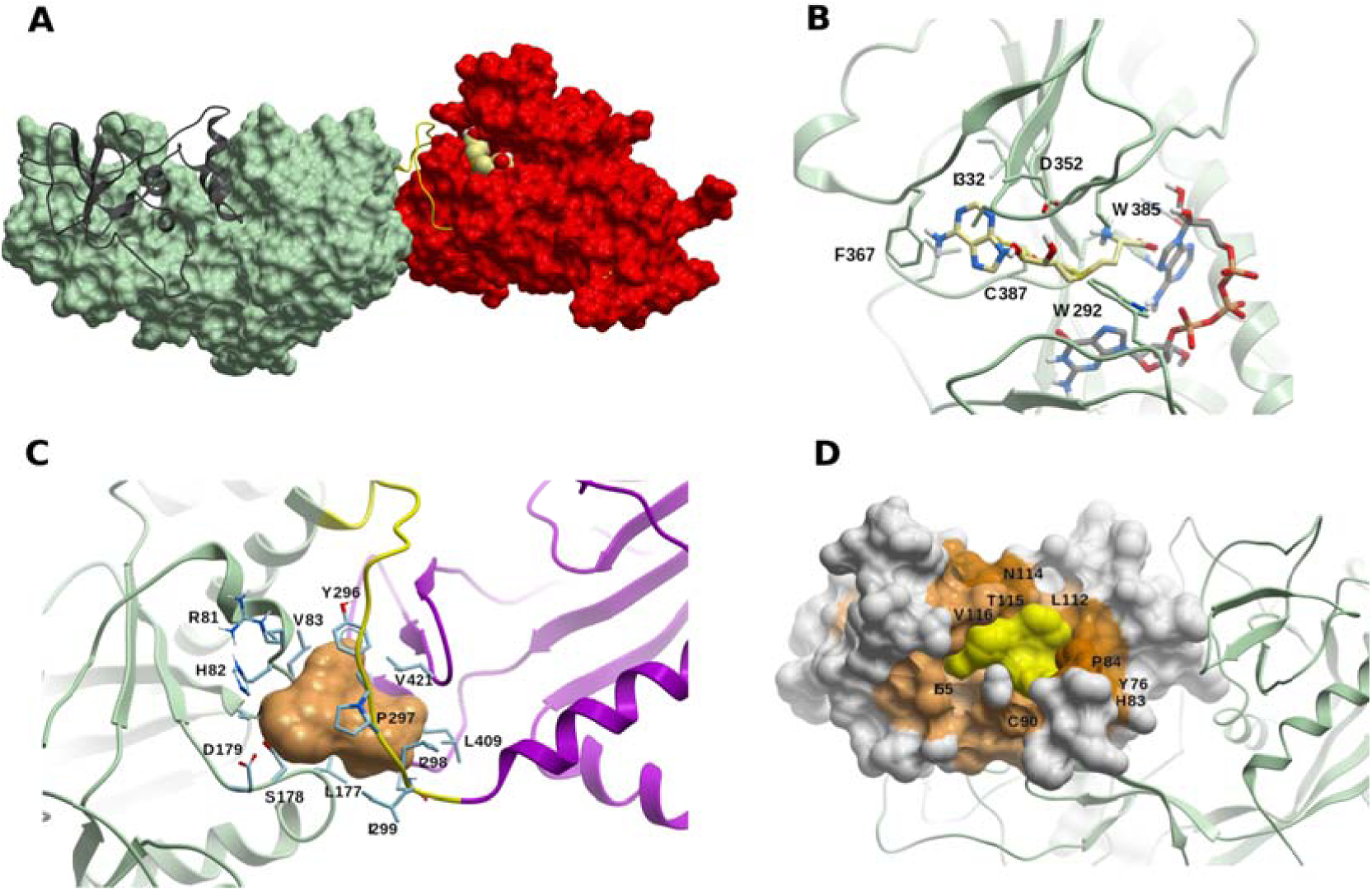
The ExoN/MTase complex (nsp14-nsp10) (A) The ExoN nsp14 domain (in green) and the MTase domain (in red) are connected by the hinge loop F286-G300 (yellow). Nsp10 is shown in dark gray ribbon, and a S-adenosyl-L-methionine (SAM) molecule (light yellow carbons) is displayed within the catalytic site. (B) A model of SAM (light yellow carbons) within the catalytic site of the MTase domain of SARS-CoV-2 nsp14. A molecule of guanosine-P3-adenosine-5′,5′-triphosphate (GpppA) (green carbons) is added as reference. (C) Potential druggable (allosteric) binding site in the vicinity of the hinge region F286-G300 (in yellow), including Y296 and P297. The linked nsp14 domains ExoN and MTase are displayed in green and magenta, respectively. (D) Druggable site (yellow surface) within a cryptic site on nsp10 (colored in brown, lighter or darker according to the cryptic score). The ExoN domain of nsp14 is shown as green ribbon.

As it has been pointed out in SARS-CoV [107,111], nsp14 has an atypical MTase fold, including also an additional α-helix in the last 12 amino acids which stabilizes the neighboring environment; deletion of this helix has been shown to decrease or abolish the MTase activity of nsp14. A third zinc finger is also present in this domain formed by C452, C477, C484 and H487, but distant from the active site, and mutations of amino acids from the zinc finger 3 have a very marginal effect on MTase activity [111], while it has been hypothesized that it might be important in binding with the nsp16-nsp10 dimer to accomplish the second methylation for completion of the capped structure. The SARS-CoV-2 N7-MTase domain in complex with methyl donor SAM is shown in Figure 9b; the binding site residues are fully conserved, thus the contacts are similar to the SARS-CoV nsp14-SAM complex. Methyl acceptor GpppA binds close to methyl donor SAM (cf. PDB 5C8S) to facilitate methyl transfer. Comparing the N7-MTase catalytic sites of SARS-CoV nsp14 structures bound to SAM (5C8T), and to SAH-GpppA (5C8S), no significant structural changes are found, thus supporting the hypothesis that ligand binding sites are pre-formed.

The hinge region (amino acids F286-G300, Figure 9a) is conserved across CoVs, thus suggesting it might have a functional role. In SARS-CoV, lateral and rotational movements of the C-terminal domain relative to the N-terminal domain of up to 13 Å have been observed [107]; moreover, from crystallographic and SAXS data it has been shown that nsp14 undergoes important conformational changes, and the hinge might act as a molecular switch. In fact, nsp14 is involved in two processes that use RNA substrates in different ways: the new synthesized RNA strand with a mismatch should be translocated by the polymerase complex to the ExoN catalytic site of nsp14, while during replication, the 5’-mRNA should go into the catalytic tunnel of the N7-MTase for methyl capping. These results could be understood considering nsp14 flexibility; moreover, hinge residues Y296 and P297 are essential for ExoN activity in SARS-CoV. The mutual dependence of the ExoN domain on N7-MTase function has been shown using mutagenesis analysis [112], and of the N7-MTase domain on ExoN activity [107]. Ferron et al. further showed that both the ExoN domain (excluding the first 71 residues) and the N7-MTase domain interact with nsp12-RdRp [107], a biologically relevant interaction due the possibility of a tandem associated activities that makes RNA replication more efficient.

Considering the structural and functional features of the SARS-CoV-2 nsp10-nsp14 dimer, the following small-molecule targeting strategies could be suggested: i) The SAM binding site is an attractive target to develop CoVs inhibitors using small-molecules that could preclude SAM or GpppA binding, thus suppressing the N7-MTase activity of nsp14. In fact, the N7-MTase catalytic tunnel lies close to the SAM binding site, and blocking it would also preclude mRNA binding; ii) A potential druggable (allosteric) site was identified using FTMap and ICM Pocket Finder defined by residues R81-A85, L177-D179, Y296-I299, N408, L409, L411, and V421. This site lies in the vicinity of hinge residues Y296 and P297 (Figure 9c). Binding of a small-molecule to this site could interfere with the dynamic behavior of nsp14 and its associated conformational changes. iii) There is a small hot spot in nsp14 (lined up by residues F60, M62, L192, M195, and K196) in the vicinity of a cryptic site, in which a molecule binding in this region would clash with nsp10 binding; iv) Considering the high sequence identity between the SARS-CoV and SARS-CoV-2 nsp10-nsp14 dimers, and that the contact residues are fully conserved, blocking the nsp10-nsp14 interaction to decrease or abolish full ExoN activity could be a valid strategy against CoV diseases. A potential borderline druggable site on the PPI surface of nsp10 with nsp14 and lined up by residues T5, E6, N40, A71, S72, C77, R78, H80, L92, K95, Y96 could be identified using FTMap (this site is defined by two hot-spots separated by ∼9 Å, what would impose a limit on the expected affinity of a potential ligand [31]. However, it should also be taken into consideration that in the absence of nsp10, the nsp14 ExoN catalytic site partially collapses and the activity decreases [113]; thus, it would be interesting to further explore the behavior of this site using MD simulations; v) A druggable site was identified within a region of high cryptic site score in nsp10 (Figure 9d). This site does not overlap with the PPI interface of nsp14 nor nsp16, and while its functional role is uncertain, it may have allosteric modulation potential.

### RNA nucleoside-2’O-methyltransferase complex (nsp16-nsp10)

Non-structural protein 16 possesses a SAM-dependent RNA 2’O-MTase activity that is capable of cap-1 formation. Nsp16 adds a methyl group to the previously capped m7GpppN by nsp14 (cap-0) to form m7GpppNm (cap-1), by methylating the ribose of the first nucleotide at position 2’-O (Figure 4). Like nsp14, nsp16 uses SAM as methyl donor [114]. In SARS-CoV, nsp16 requires nsp10 to execute its activity, since nsp10 is necessary for nsp16 to bind both m7GpppA-RNA and SAM [115]; moreover, nsp16 was found to be an unstable protein in isolation [116], and most of the disruptions in the interface of nsp16-nsp10 abrogate the methylation activity [117]. In humans, most mRNAs include the cap-1 modification; while cap-0 appears to be sufficient to recruit the entire translational machinery, this modification is necessary to evade recognition by host RNA sensors, such as Retinoic acid-Inducible Gene I (RIG-I), MDA-5, and interferon induced proteins with tetratricopeptide repeats (IFIT), and to resist the IFN-mediated antiviral response [118].

Several crystal structures of the nsp16-nsp10 heterodimer have been recently solved (cf. Table S4) (Figure 10a). SARS-CoV-2 nsp10 is 99% identical to SARS-CoV, while 59% identical to MERS-CoV. Similarly, nsp16 is highly similar to SARS-CoV (95%), but only 66% identical to MERS-CoV. Both proteins interact through a large network of hydrogen bonds, water-mediated interactions, and hydrophobic contacts [119]. The high conservation of nsp10 and nsp16 sequences, and the complete conservation of catalytic and substrate-binding residues strongly support the idea that nsp16-mediated 2’-O-MTase mechanism and functionality are highly conserved in CoVs. Nsp10 exhibits two zinc fingers: the first one is coordinated by C74, C77, H83 and C90, and the second one is coordinated by C117, C120, C128, and C130. The zinc fingers residues are 100% conserved across β-CoVs, highlighting the relevance of this motif for nsp10 activity in the replication process.

**Figure 10.**
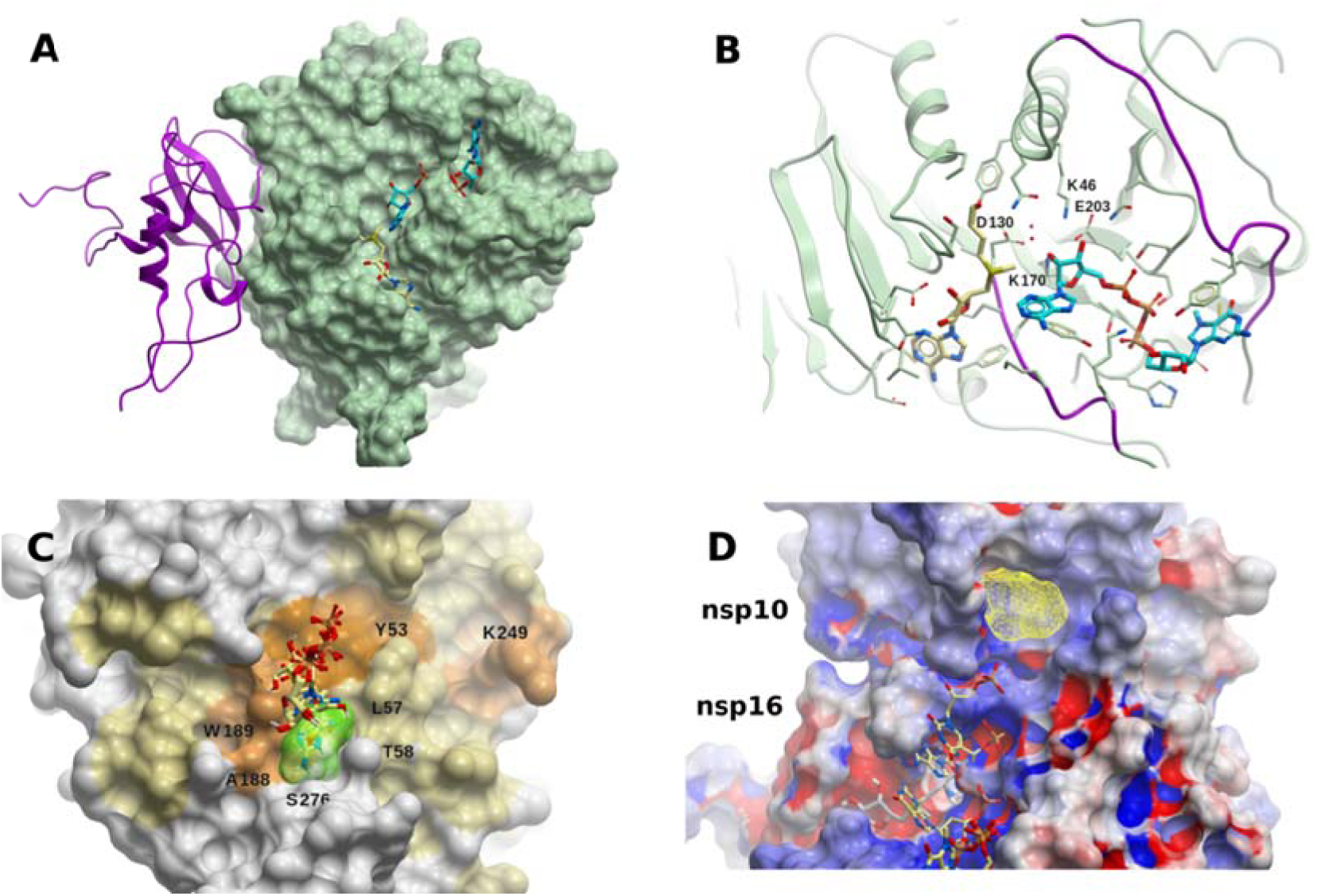
The SARS-CoV-2 RNA nucleoside-2’O-methyltransferase complex (nsp16-nsp10). (A) Structure of the 2’O-MTase (nsp16, green molecular surface) heterodimer with nsp10 (magenta ribbon). Molecules of SAM (light yellow carbons) and m7GpppA (cyan carbons) are shown within the catalytic site. (B) Catalytic site of nsp16 with methyl donor SAM (light yellow carbons) and methyl acceptor m7GpppA (cyan carbons). The nucleotide binding site flexible loops (D26-K38, M131-N138) are colored in blue. The highly conserved KDKE motif (K46, D130, K170, E203) for methyl-transfer, found in many 2’O-MTases, is highlighted in magenta, and oxygen water molecules are displayed in red. (C) Binding hot spot (transparent yellow) identified with FTMap in the vicinity of residues L57, T58, A188, C209, N210, and S276, on the surface of nsp16, within an extended cryptic site identified in the same region using CryptoSite (brown colored surface). Small-molecules bound within that site (taken from crystal structures) are also displayed (light yellow carbon atoms): [adenosine, 2-(n-morpholino)-ethanesulfonic acid, β-D-fructopyranose, 7-methyl-guanosine-5’-triphosphate, and 7-methyl-guanosine-5’-diphosphate]. This site lies on the opposite side of the catalytic site, ∼25 Å away from it, in what thus could be an allosteric site. (D) Extension of the RNA groove in nsp16 towards nsp10. Five RNA nucleotides are shown (light yellow carbon atoms), which correspond to those of the human mRNA 2’O-MTase (PDB 4N48), after structural superposition of the binding site residues. SAM is displayed in grey carbon atoms. The consensus site identified with FTMap is shown in yellow mesh. Nsp16 and nsp10 are colored according to their electrostatic potential (blue, positively charged; red, negatively charged; white, neutral).

Most nsp16-nsp10 structures feature SAM (or the close analog pan-MTase inhibitor sinefungin) in its substrate binding site, being coordinated by residues N43, Y47, G71, A72, S74, G81, D99, N101, L100, D114, and M131, together with few water molecules (Figure 10b). All these residues are also 100% conserved in SARS-CoV. Some structures also include m7GpppA within the catalytic site (PDBs 6WQ3 and 6WRZ in presence of SAH, and 6WKS and 6WVN in presence of SAM). The nucleotide binding site is surrounded by flexible loops comprised of amino acids D26-K38, M131-N138, and the highly conserved KDKE motif (K46, D130, K170, E203) for methyl-transfer, which is found in many 2’-O-MTases (including the phylogenetically more distant MERS-CoV), is also present (Figure 10b). The MTase active site is clearly an attractive target for antiviral drug discovery, but the structural conservation among CoVs and host cellular MTases might pose a challenge for the development of a specific compound. Several inhibitors for various MTases were developed from sinefungin analogs [120].

Using FTMap, a small hot spot was found in the vicinity of residues L57, T58, A188, C209, N210, and S276, within an extended cryptic site on the surface of nsp16 (Figure 10c). Interestingly, recent crystal structures of the nsp16-nsp10 dimer feature small-molecules bound within that site, such as adenosine (PDB 6WKS), 2-(n-morpholino)-ethanesulfonic acid (PDB 6YZ1), β-D-fructopyranose (PDB 6W4H), 7-methyl-guanosine-5’-triphosphate (PDBs 6WVN and 6WRZ), and 7-methyl-guanosine-5’-diphosphate (PDB 6WQ3) (Figure 10c). This site lies on the opposite side of the catalytic site, 25 Å away from it, in what could thus be an allosteric site [121]. Further studies are necessary to clarify the function of this binding site, and its impact and feasibility in terms of druggability.

Considering that m7GpppAC_5_ acts as an effective substrate of SARS-CoV nsp16-nsp10 [122], Chen et al. performed docking of m7GpppGAAAAA and m7GpppAAAAAA within the SARS-CoV nsp16-nsp10 dimer SAM [115], and found that while the first three nucleotides contact nsp16, the remaining ones are in contact with nsp10, which extends the positively charged area of the RNA-binding groove. Interestingly, all the residues defining this groove extension are also conserved in SARS-CoV-2. The extension to the RNA-binding site provided by nsp10 may serve to accommodate the RNA chain and stabilize the interaction between m7GpppA-RNA within the catalytic site, what could also be observed superposing the SAM binding site of nsp16 with that of human mRNA 2’O-MTase (PDB 4N48) [119] (Figure 10d). Using FTMap, we identified a small borderline druggable site within nsp10, in the area where RNA would extend (Figure 10d). This pocket lies within the RNA binding groove extension described above, and is lined up by the side chains of amino acids P37, T39, C41, K43, F68, A104, and P107 of nsp10. A molecule binding to it might interfere with extended RNA binding. It should be stressed that, due to its zinc fingers, nsp10 has also the ability to bind poly-nucleotides non-specifically [123].

It was also shown that 12-mer and 29-mer peptides extracted from the dimerization surface of SARS-CoV nsp10 (F68-H80 and F68-Y96, respectively) were found to inhibit the activity of nsp16 at IC_50_ ∼ 160 μM [124]; this suggests that using short peptides might be a possible strategy, considering there is 100% sequence conservation within the nsp10/nsp16 interface for SARS-CoV and SARS-CoV-2. Interfering with the nsp16-nsp10 dimer formation using small molecules might still be challenging, however, due to the large area of contact and the absence of buried pockets at the the nsp10-nsp16 interaction surfaces. In spite of this, a cryptic site was identified on nsp16 in the region of the PPI interface with nsp10 (Figure S5), close to where the 12-mer and 29-mer peptides would bind. However, no CSs were found near this cryptic site, so further studies should be performed to confirm this site as a potential target to inhibit dimerization.

### RNA uridylate-specific endoribonuclease (nsp15)

The nsp15 harbors a nidoviral RNA uridylate-specific endoribonuclease (NendoU) that belongs to the EndoU family, whose activity is to cleave downstream of uridylate, releasing 2’-3’ cyclic phosphodiester, and 5′-hydroxyl termini. The NendoU cleaves polyuridines from viral ssRNA, produced during the priming of the poly(A) ssRNA in replication (Figure 3), which helps to dampens dsRNA melanoma differentiation-associated protein 5 (MDA5)-dependent antiviral IFNresponses [125], thus evading the host response. Clearly, small molecules that block the RNA catalytic sites, or interfere with the oligomeric formation of nsp15, would inhibit the enzyme and help trigger cellular antiviral mechanisms.

The SARS-CoV-2 nsp15 structure displays an N-term oligomerization domain, a middle domain, and a C-term NendoU catalytic domain [126] (Figure S6a). Nsp15 has been crystallized in its apo form (PDB 6VWW at 2.2 Å resolution), and in complex with citric acid (PDBs 6XDH at 2.35 Å, and 6W01 at 1.9), uridine-3’-monophosphate (PDB 6×4I at 1.85 Å), uridine-5’-monophosphate (PDB 6WLC at 1.85 Å), drug Tipiracil (PDB 6WXC at 1.85 Å), and product di-nucleotide GpU (PDB 6×1B at 1.97 Å). All structures are very similar, with main chain RMSD values of less than 0.5 Å between any pair of them. Taking structure 6W01 as a reference, SARS-CoV-2 nsp15 exhibits ∼0.5 Å RMSD with respect to its SARS-CoV counterpart (88% sequence identity, 95% similarity), and 1.1 Å with respect to the nsp15 from MERS-CoV (51% sequence identity, 65% similarity) (cf. Ref. [126] for a detailed comparison of SARS-CoV-2 nsp15 with SARS-CoV and MERS-CoV structures).

The catalytic active site within the NendoU domain is formed by residues H235, H250, K290, V292, S294, T341, and Y343 (Figure S6b) (also conserved in SARS-CoV and MERS-CoV nsp15s), with H235, H250, and K290 as the proposed catalytic triad [126]. The catalytic residues also form a druggable binding site, as can be confirmed by its shape and the crystal structure of drug Tipiracil in complex with nsp15 (PDB 6WXC). The catalytic activity has been observed to be metal-dependent in most NendoU proteins, although there could be exceptions in other nidoviruses [127]. Although no crystal structure has been found with a metal ion (nor complexed with RNA), a metal binding site required for maintaining the conformation of the active site and substrate during catalysis has been proposed to be coordinated by the carboxylate of D283, the hydroxyl group of S262, and the carbonyl oxygen of P263.

SARS-CoV-2 nsp15 folds into a hexamer (a dimer of trimers) [126], in agreement with an earlier work showing that SARS-CoV nsp15 conformationally exists as a hexamer (PDB 2RHB, [128]), and also suggesting an oligomerization-dependent endoribonuclease activity [129]. The channel of the hexamer is ∼10 Å wide, open from top to bottom, and the hexamer is stabilized by extensive contacts between monomers, and as such, it might be disrupted or destabilized by mutations or small molecules [126]. Using FTMap, we identified two potential druggable sites; the first one is delimited by residues K71, N74, N75, M272, S274-N278, I328, L346, and Q347 (Figure S6c), and close to the binding area of another protomer within the hexamer (Figure S6c); a small-molecule binding to this site might disrupt PPI; the second one, delimited by residues E69, K71, K90, T196, S198, L252, D273, K277, Y279, V295, and D297 is deep and rather buried, within an area with higher-than-average cryptic site score and an opening towards the hexamer channel.

### Non-structural protein 9 (nsp9)

Nsp9 acts as a hub that interacts with several viral components, binds ssRNA, nsp8, the N protein, and several host components as nuclear pore proteins [15,130,131]. Proteomic experiments showed that SARS-CoV-2 nsp9 also binds to several nuclear pore proteins [15], a process well characterized for other viruses that affect nuclear shuttling as part of their replicative and host shutdown mechanisms [132]. SARS-CoV nsp9 exists in solution as a homodimer through the interaction of the GXXXG motif in opposite parallel α-helices of the protomers [130]; mutations in any of the glycines inducing dimer disruption have been associated with impeded viral replication [133,134], and reduced RNA binding in SARS-CoV [134] and in porcine delta coronavirus (PDCoV) [135]. Thus, as it has been already suggested, disruption of the homodimeric interface could be an appealing strategy against CoV-associated diseases [136]. Homologs of nsp9 have been found in other β-coronaviruses such as SARS-CoV, MERS-CoV, HCoV-229E, and the avian infectious bronchitis virus, with different degrees of sequence similarity compared to SARS-CoV-2 nsp9. The mechanism of RNA binding to nsp9, and how it enhances viral replication, are not yet fully understood, but apparently depends on several positively charged aminoacids on the surface [135]. While it is not known whether SARS-CoV-2 nsp9 plays an analogue role as its SARS-CoV counterpart, their high sequence identity (97%; they differ in only three amino acids out of 113), and their strong structural similarity (see below), might suggest a strong conserved functional role.

The structure of the SARS-CoV-2 nsp9 homodimer has been solved at 2.0 Å (PDB 6WXD) [137], and also at 3.0 Å (PDB 6W4B); both structures share a backbone RMSD of 0.5 Å and 0.9 Å for the monomer and dimer, respectively. The monomer has a backbone RMSD of 0.94 Å compared to that of SARS-CoV (PDB 1QZ8, [138]); the RMSD value decreases to 0.44 Å when considering α-helices and β-sheets only. The protomers interact mainly through van der Waals contacts of the backbone from the conserved GXXXG motifs within the opposed α-helices, near the C-terminus of the protein (Figure S7a), in a similar fashion to nsp9 of other CoVs [136,138,139]. Another nsp9 structure was determined including in the N-terminal tag a rhinoviral 3C protease sequence LEVLFQGP (PDB 6W9Q) [137]. The inserted peptide folds back and forms a β-sheet with the N-terminal of the protein. Residues LEVLF of the peptide make hydrophobic contacts with residues P6, V7, A8, L9, Y31, M101, S105, and L106 of nsp9, and is hydrogen bonded with residues P6, V7, L9 and S105. These nsp9 residue are conserved among other nsp9 homologs, including 100% conservation in SARS-CoV. This provides evidence that this binding site could be targeted by a peptidomimetic small molecule to disrupt dimer formation and thus reduce RNA binding and viral replication.

Using FTMap, two potential druggable sites were identified in nsp9. One site is defined by residues R39-V41, F56-S59, I65-E68, I91, and K92, the other one by residues S13, C14, D26-L29, L45-L48, and K86 (Figure S7b). These two sites lie in regions of intermediate-to-high cryptic site score, and do not overlap with the dimerization interface, but are close to positively charged residues that might be involved in RNA binding, as was postulated for IBV [140]; it should also be considered they might interfere in nsp9-nsp8 PPI, though this is subject of validation. A hot spot within a region of moderate-high cryptic site score was identified in the C-terminus of nsp9, delimited by residues C73, F75, L88, L103, L106, A107, and L112 (Figure S7c); a molecule binding in this region might clash with the N-terminal part of the other protomer.

### ADP-ribose-phosphatase (nsp3 domain)

The first nsp3 macrodomain (Mac1) is conserved throughout CoVs and has an ADP-ribose phosphatase (ADRP) activity, by which ADRP binds to and hydrolyzes mono-ADP-ribose [141]. This appears to be related to the removal of ADP-ribose from ADP-ribosylated proteins or nucleic acids (RNA and DNA); it should be pointed out that ADRP is able to remove mono-ADP-ribose, but not poly-ADP-ribose [141]. Anti-viral ADP-ribosylation is a host post-translational modification in response to viral infections, since many of the IFN and cytokine signaling components, as NF-kappa-B essential modulator (NEMO), TANK-binding kinase 1 (TBK1), NFκB among others need to be ribosilated to be fully active [142,143]. Although ADRP is not an essential protein for viral replication, it has been shown to be an essential pathogenesis factor in animal models for CoV infection; for example, mutations of ADRP in SARS-CoV enhanced IFN response and reduced viral loads *in vivo* in mice models [144,145]. Thus, its role against host-induced anti-viral activity makes it an attractive target for drug design.

Recently, five crystal structures of SARS-CoV-2 ADRP were solved (see Table S1) [146], including the apo form (PDBs 6VXS at resolution 2.0 Å, 6WEN at 1.35 Å), and in complex with 2-(N-morpholino)ethanesulfonic acid (MES) (PDB 6WCF at 1.09 Å), AMP (PDB 6W6Y at 1.45 Å), and ADP-ribose (PDB 6W02 at 1.50 Å) (Figure S8a). Another structure of ADRP complexed with ADP-ribose (PDB 6WOJ) at 2.2 Å is available. These structures exhibit low main chain atom RMSD between any pair of them, with values in the range 0.25-0.65 Å. The SARS-CoV-2 ADP-bound ADRP structures share structural similarity to related homologs from SARS-CoV (71% sequence identity, 82% similarity) and MERS-CoV (40% sequence identity, 61% similarity), with main chain RMSD values of 0.6 Å (PDB 2FAV) and 1.4 Å (PDB 5HOL), respectively.

The binding site of SARS-CoV-2 ADRP (Figure S8b) bears high similarity with those of SARS-CoV and MERS-CoV. The ADP-ribose is stabilized within the binding site through hydrophobic interactions, and direct and solvent-mediated hydrogen bond interactions; it should be highlighted that most of the hydrogen bonds involve main chain atoms. The binding mode of ADP-ribose is conserved in those three CoV (RMSD values of 0.3 Å and 1.2 Å, respectively, with respect to SARS-CoV-2 ADRP), with ADP-ribose exhibiting similar affinities towards the three CoVs [141]. Within the overall conserved binding site conformation of the SARS-CoV-2 ADRP structures, some shifts are observed comparing the apo structure, and those in complex with AMP, ADP-ribose, and MES, mainly around the proximal ribose. The rotameric states of several side chains (F132, I131, F156) adopt a ligand-dependent conformation, and a flip in the A129-G130 peptide bond could be observed, dependent on the presence of the phosphate group in ADP-ribose (phosphate 2), or MES.

Considering the highly conserved structural features CoVs, especially SARS-CoV and MERS-CoV, and its role in countering host-induced antiviral responses, ADRP appears as an attractive therapeutic target. It should be noted that no other druggable binding site could be identified other than the ADP-ribose pocket using FTMap or ICM Pocket Finder, nor could cryptic sites be found on the surface.

### Ubiquitin-like 1 domain (nsp3)

The first ∼110 residues of nsp3 have an ubiquitin-like fold, and are thus named the Ubl1 domain. The function of Ubl1 in CoVs is related to ssRNA binding, while probably also interacting with the N protein. In SARS-CoV Ubl1 has been shown to bind ssRNA with AUA patterns, and since the SARS-CoV 5’-UTR (un-translated region) is rich in AUA repeats, it is likely it binds to it [147]; in fact, SARS-CoV-2 has 439 AUA within its genome. In MHV, this Ubl1 domain binds the structural N protein [148,149]. Both putative activities point to track the viral RNA to DMVs, based in the nsp3 localization in DMVs and the Ubl1 binding to RNA and the N protein. It was also shown that MHV-Ubl1 is essential for viral replication, since viable virus could not be recovered from the Ubl1 full deletion mutant [148].

There is yet no available experimentally solved structure of SARS-CoV-2 Ubl1. Considering that the sequence identity and similarity with its SARS-CoV counterpart is 79% and 93%, respectively, (with only one deletion in the alignment, close to the N-terminal), a structural model was built using SARS-CoV Ubl1 (PDB 2GRI) as template (Figure S9a). There are several positive residues on the protein surface, compatible with ssRNA binding. These residues are conserved in SARS-CoV, with the exception of R23N, R102H, and N98K (SARS-CoV numbering). Two small potentially druggable sites were identified with FTMap and ICM Pocket Finder, which lie in areas of above-average cryptic site score. One site is defined by the side chains of Y42, T43, T48, E52, F53, C55 and V56 (Figure S9b), and the second one by F25-D28, T86, Y87, W82, and C104-F106 (Figure S9c). These sites are near positively charged residues, but do not overlap with them. While a molecule binding to them might interfere with ssRNA binding, it is also possible that it might disrupt PPIs with partner proteins, such as the N protein. Further biochemical and functional characterization of Ubl1 is needed to shed light on the actual value of these sites.

### SARS-unique domain (SUD) (nsp3 domain)

The SUDs binds ssRNA with different base affinities [150], and as a part of nsp3, is implicated in tracking the viral RNA to DMVs, a subcellular space where viral replication machinery is concentrated. In SARS-CoV-2 and SARS-CoV there are three SUD domains connected by short peptide linkers, SUD-N, SUD-M, and SUD-C, indicating the N-terminal, the middle, and the C-terminal regions of SUD. SUD-N (macromolecular domain 2, Mac2) binds G-quadruplexes, an unusual nucleic-acid structures formed by guanidine-rich nucleotides in ssRNAs; it shares 71% of identity and 85% of similarity with its SARS-CoV counterpart. SUD-M/Mac3 (macromolecular domain 3) has a single-stranded purine rich (G and/or A) RNA binding activity [151] which includes poly(A) [151] and G-quadruplexes [152]; this allows SUD-M to act as a poly(A)-binding-protein and thus protect the poly(A) tail, at the 3 ‘end of viral RNA, from 3’ exonucleases. SUD-C/DPUP (Domain Preceding Ubl2 and PL^pro^) also binds to ssRNA, and recognizes purine bases more strongly than pyrimidine bases [151]. This RNA binding activity is apparently stabilized by the presence of SUD-M, while SUD-C seems to modulate the sequence specificity of SUD-M [151]. Mutagenesis analysis in SARS-CoV showed that SUD-M is indispensable to the virus replication, while the absence of SUD-N or SUD-C barely reduce the virus titer [153].

The structure of the SARS-CoV construct SUD-N/SUD-M (SUD-NM) has been solved by crystallography at 2.2 Å resolution (PDB 2W2G). The solution structures of SARS-CoV SUD-M and SUD-C within a SUD-MC construct were obtained using NMR (PDBs 2KQV and 2KQW, respectively), together with the isolated SUD-C (PDB 2KAF). The isolated SARS-CoV SUD-M has also been solved by NMR (PDBs 2JZD and 2JZE), together with the structure of an N-terminal extended SUD-M (PDBs 2RNK and 2JZF). The main chain RMSD values among the corresponding solved structures is within 0.8 Å. It has been shown that SUD-NM is monomeric in solution [151], and the absence of evidence suggesting a tight transient or static contacts in solution suggested that SUD could be modeled as three flexible linked globular domains [151].

The SARS-CoV-2 SUD domains are very similar to their SARS-CoV counterpart, with sequence identity (similarity) of 71% (91%), 79% (94%), and 72% (92%) for SUD-N, SUD-M, and SUD-C, respectively. Considering that only SUD-M appears to be essential to viral replication, we focus our druggability analysis on it. We thus built a model by homology using PDB 2AKF as template. For SARS-CoV, it was shown that upon poly(A) binding to SUD-MC, the molecular surface area of SUD-M affected by the NMR chemical shift perturbation experiments was mapped to a positively charged surface cavity [151], defined by residues N532, L533, I556, M557, A558, T559, Q561, and V611 [154]. This area is wholy conserved in SARS-CoV-2. A potential druggable binding site was identified on this area using FTMap and ICM Pocket Finder (Figure S10), and we could hypothesize that a small-molecule binding to this site might preclude RNA binding.

### Nucleic-acid binding region (NAB, nsp3 domain)

NAB is a small domain of ∼120 amino acids that binds ssRNA, strongly preferring sequences containing repeats of three consecutive guanosines [155]. In SARS-CoV NAB binds ssRNA through a positively charged surface patch defined by the residues K75, K76, K99, and R106, while the neighboring residues N17, A18, S19, D66, H69, T97 are also affected by RNA binding [155]. Interestingly, this RNA binding site bares similarity to that of the sterile alpha motif of the Saccharomyces cerevisiae Vts1p protein [155]. Experiments have shown that N-terminal and C-terminal extensions, corresponding to links with the PL^pro^ and the TM-Lumen/Ectodomain, respectively, behave as flexibly disordered segments [155].

While the SARS-CoV-2 NAB structure has not been solved yet, the SARS-CoV counterpart has been solved by NMR (PDB 2K87 [155]). We used the latter to build a structural model of SARS-CoV-2 NAB by homology (sequence identity 81%, similarity 94%, no gaps in the alignment). All the positive residues are conserved, with two additions, T43K and S51K (Figure S11a).

No druggable binding sites were identified within the positive charged patch on the surface where ssRNA binds in SARS-CoV; however, two cryptic sites with nearby CSs were predicted on the sides of that patch (Figure S11b). It should be further explored whether these sites are involved in PPI, and whether small-molecules binding to these sites might allosterically modulate ssRNA binding.

### Other nsp3 domains

The DUF3655/HVR/Acidic-domain, TMs-Lumen/Ectodomain, and the C-Terminal domain/Y-Domain (Table S1) are structurally uncharacterized nsp3 domains, which bear little similarity to any other experimentally solved structure, thus ruling out the possibility of homology modeling.

DUF3655-HVR is a hypervariable, Glu and Asp rich domain, probably implicated in protein-protein interactions with N, as in MHV [156]. Since is not essential to viral replication in MHV [148], it would be a second priority for drug targeting.

The Lumen/Ectodomain is flanked by two transmembrane domains (TMs), exposed to the ER lumen, and works by binding to the lumen domain of nsp4, necessary to form the Double-Membrane Vesicles (DMVs) where CoVs replicate, anchoring the whole nsp3 protein to membranes through the TMs. While DMVs formation is essential, the lack of structural information poses an insurmountable hurdle for drug discovery.

The C-Terminal domain/Y-Domain is conserved in CoVs, and it seems to be involved in the DMVs formation, probably interacting with nsp6, and improving nsp3-nsp4 interaction [157]. Although breaking this interaction would seriously impact on viral replication, the lack of structural and biochemical information precludes any targeting attempt.

### Other non-structural proteins: nsp1, nsp2, nsp4, and nsp6

At this point is clear that, both structural and replicative core proteins are essential to the virus. Deletional studies in SARS-CoV and MHV have shown that nsp1 and nsp2 are not essential for viral replication, but their absence might affect the final viral titer [158,159], while nsp4 and nsp6 seems to be necessary for Double-Membrane Vesicles (DMVs) formation.

Nsp 1 is the leader protein and the first translated and PL^pro^-processed protein. It has the capacity to bind to the 40S ribosome subunit to inactivate the translation of host mRNAs [160], also selectively promoting host mRNAs degradation, which makes it the main actor in host cell shutdown [160], and indirectly in the immune response evasion [161] by delaying IFN responses [162]. Additionally, based on recent proteomics analysis, SARS-CoV-2 nsp1 seems to interfere with the host DNA duplication by interacting with the DNAPolA complex, that is the host primase complex, the first step in DNA synthesis [15]. Although nsp1 is not essential for viral replication, its absence makes the virus susceptible to IFN [163], what would make it an important pathogenic factor and a good target for drug design. However, the structural information available of this 180-amino acids protein is limited. There is an NMR structure from SARS-CoV (PDB 2GDT) of 116 amino acids (corresponding to H13 to G127 in SARS-CoV-2), while the active sites for RNA and 40S binding involve residues R124 and K125, and K164 and H165, respectively [164]. There is a recent cryo-EM structure of the nsp1-C-terminus (E148-G180) in complex with the 40S ribosomal subunit and RNA (PDB 6ZLW [160]). While amino-acids K164 and H165 are present, the C-terminal portion of only 33 residues cannot be used for structure-based drug design.

Nsp2 is a membrane protein not essential for the viral production in SARS-CoV and MHV homologs, however, in MHV, the presence of nsp2 positively affects the viral titer [165]. It is a delocalized membrane protein, but is recruited to replication sites by other viral components [158], probably by M [166]. It has a low sequence conservation across CoVs, which could imply that its function cannot be conserved among different species, which supports its not being essential to viral function [167]. In SARS-CoV-2, proteomic data showed that nsp2 interacts with cellular components associated with vesicles formation and translational regulators [94], which allows us to infer that its function is associated with the formation of membrane structures, and co-opting host components. However, up to the present date there is no structural data of nsp2 from SARS-CoV-2 or related proteins.

Nsp4 is essential for viral replication in MHC [168]. The main function of nsp4 is the formation of Double-Membrane Vesicles (DMVs) in SARS and MERS [40]; DMVs are very important since they concentrate the viral replication machinery. In SARS-CoV and MERS-CoV, nsp4 has four TM domains, a ER-Lumen exposed domain, and a cytoplasmatic exposed C-Terminal domain [169]. In SARS and MHV, the lumen domain of nsp4 interacts strongly with the nsp3 lumen domain/ectodomain to induce DMVs formation, DMVs are where the viral replication machinery is recruited and the viral replication happens [157]. Clearly, a nsp3-nsp4 structure would allow to study the design of a PPI inhibitor that could prevent DMVs formation. However, only the cytoplasmatic C-Terminal domain (∼90 amino acids) of MHV and Feline CoV nsp4 structures have been experimentally solved (3VC8, 3GZF, respectively). With SARS-CoV-2 nsp4, these proteins share 59% and 39% identity, and 76% and 55% similarity, respectively [170,171].

Nsp6 is a membrane protein with a medium conservation degree between different CoVs. In SARS-CoV, nsp6 collaborates with nsp4 and nsp3 in DMVs formation, also inducing the formation of membrane vesicles [40]. Additionally, in several CoVs, nsp6 activate omegasome and autophagosome formation independently of starvation, to degradate cellular components to increase the availability of resources for viral replication [172]. Nevertheless, there is no structural information of any nsp6 related protein which would allow structure-based drug discovery.

### S protein

The SARS-CoV-2 spike-protein (S protein, ORF2) is a large homotrimeric multidomain glycoprotein. The small C-terminal transmembrane attaching domain is followed by the S2 and S1 subunits. During infection, the receptor binding domain (RBD) in S1 is exposed and recognized by ACE2, the junction S1/S2 is cleaved by a furin-like protease releasing the S1 domain, while S2 is cleaved again, by the metalloprotease TMPRSS2, to expose the fusion peptide (FP), which is responsible to induce the membrane fusion mechanisms [25,173] (Figure 1).

Several strategies are focused on inhibiting the viral entry, either by interfering with the binding of S to ACE2, or with the membrane fusion induction [20]. The first approach was the development of inhibitory antibodies or recombinant ACE2 proteins that block the S protein [20]; as the outer surface component of CoVs, the S protein is a major target of antibodies, and the main focus of vaccine development. Using a different approach, based on structural information and the knowledge of the membrane fusion mechanisms, several peptides that mimic neutralizing antibodies have been developed [20]; for example, a series of lipo-peptides EK1C1-EK1C7, and IPB02, that targeted the subdomain HR1 in the S2 fragment to inhibit membrane fusion by interfering with the FP, and which showed a decrease of SARS-CoV-2 virus titer in cell cultures [174,175].

As the mechanism of the virus entry depends on host factors, targeting ACE2 and TMPRSS2 is being explored. There are at the least three ACE2-inhibitors (Captopril, Lisinopril, Losartan), but this option is not very appealing due to blood pressure decreasing side-effects. The inhibition of TMPRSS2 is also being explored, by blocking the proteolytic priming of the S2 fragment, necessary to induce the membrane fusion; the inhibitors Nafamostat [176], Gabexate, and Calmostat [177], have been shown to inhibit viral production in cell cultures, and will be tested in humans; however, their low half-life opens the door to developing more efficient new drugs (cf. Ref. [178] for new options targeting TMPRSS2).

The promotion of ADAM17 activity, a metalloprotease that inactivate ACE2 by shedding, is an alternative actually approached by the use of chloroquine and its analogs, which inhibit SIGMAR1-2 receptors responsive to activate ADAM17 [179].

### E Protein

It is a small homopentameric membrane protein (75 amino-acids per protomer) [180], which has been shown to be essential for viral particle assembly in SARS-CoV [51]. The N-terminal region of the protein spans the lipid bilayer twice, while the C-terminal is exposed to the interior of the virus [181,182]. In IBV and SARS-CoV, the E protein interacts with the viral M protein through an undefined region [183,184]; and also with the host Protein Associated with Lin Seven 1 (PALS1), a factor associated to the pathogenesis [185], thought the E protein C-terminal. Also, in many CoVs, the E protein works as an ion-channelling viroporin [180], which affects the production of cytokines, and in consequence the inflammatory response [186].

While there is no structural data available for the SARS-CoV-2 E protein, the structure of the TM region of the SARS-CoV homopentamer is available (residues 8-65) (PDB 5×29). Within that region, the corresponding E proteins share 91% sequence identity, and 98% sequence similarity. The homology model of the SARS-CoV-2 E protein is shown in Figure S12. Based on structural and functional considerations, the E protein channel would be the primary target site for drug development. In fact, hexamethylene amiloride (HMA) has been reported to bind to the SARS-CoV E protein homopentamer, but not to an isolated protomer [187].

### N protein

β-coronavirus nucleocapsid (N) proteins are involved in the packing of viral +ssRNA to form a ribonucleoprotein (RNP) complex, which interacts with the M protein [49]. They share an overall conserved domain structure, with an RNA-binding N-terminal domain (NTD, amino acids 49-175), and a dimerization C-terminal domain (CTD, amino acids 247-365); these two domains are connected by a disordered region, and the C-terminal tail at the CTD (366-419) is termed the B/N3 domain. The CTD forms a homodimer in solution, while addition of the B/N3 spacer results in homotetramer formation [188]. It has been suggested that the assembly of β-coronavirus N protein filaments may consist of at least three steps, namely, dimerization through the CTD, tetramerization mediated by the B/N3 region, and further filament assembly through both viral RNA binding and association of N protein homotetramers [188]. Since the formation of the RNP complex is essential for viral replication, identification of small-molecule modulators of the nucleocapsid assembly, interfering with NTD RNA binding or precluding CTD dimerization or oligomerization, would be valid therapeutic strategies against SARS-CoV-2.

Several structures of the N protein NTD and CTD have recently become available (see Table S5). Structures of the NTD (and CTD) superimpose closely among themselves. With respect to SARS-CoV, the NTD and CTD share 88% (96%), and 96% (98%) sequence identity (similarity), respectively, and the corresponding structures overlay with RMSD values within 0.8 Å.

The structure of the NTD solved by NMR (PDB 6YI3) showing that amino-acids A50, T57, H59, R92, I94, S105, R107, R149, and Y172 participate in RNA binding [189] is shown in Figure S13a. A druggable site identified near those residues partially overlaps with the pose of AMP bound to the NTD of the N protein of the homolog protein human CoV OC43 (HCoV-OC43) (PDB 4LI4), in agreement with an earlier hypothesis [24]. Two small borderline druggable sites have been also identified with FTMap (site 1: L159-P162, T165, L167, A173, S176; site 2: Q70-N75, Q83, T135, P162). These sites would not overlap with RNA binding, and further experiments are needed to confirm whether they could be allosteric sites, or within PPI interfaces.

The CTD homodimer structure is shown in Figure S13b (PDB 6WZQ), also displaying two potential druggable sites, where a molecule binding to any of them might interfere with dimer formation. Two other distinct sites were predicted on the surface of the homodimer, one over the central four-stranded β-sheet, and the other opposite to it, near the C-terminal α-helices (Figure 13c). Since it is not clear yet how the homotetramer is formed [188], it should be further explored whether any of these sites might overlap with PPI interfaces.

### M protein

It is a membrane homodimeric glycoprotein that forms part of the virion. In SARS-CoV, the M protein interacts with the N protein, being a nexus between virus membrane formation and RNA association in the virion [49,190], also inhibiting IFN production in SARS and MERS diseases [191,192]. While it is stablished that dimerization and N-M PPI motifs reside in the C-terminus of the protein [193], the lack of structural data for SARS-CoV-2 and other CoV homologs precludes further drug discovery targeting protein-M.

### Orf3a/X1/U274

Orf3a was characterized as a potassium ion channel in SARS-CoV, involved in inducing caspase-dependent apoptosis under different pathologic conditions [194]. It is included in the virion [195], and also interacts specifically with the M, E, and S structural proteins, as well as with Orf7a/U122 [196]. In SARS-CoV, orf3a expression increases the mRNA levels of all three subunits of fibrinogen, thus promoting fibrosis, one of the serious pathogenic aspects of SARS [197], and the expression of NFκB, IL8, and JNK, all involved in inflammatory responses [198]. Thus, since orf3a is responsible for inducing two of the serious pathological affections induced by SARS-CoV, design therapeutics that suppress its function could be very important; moreover, the presence of similar proteins to orf3a other β-CoVs (SARS-CoV-2 orf3a shares ∼73% identity and ∼85% similarity with its SARS-CoV counterpart) and in α-CoVs and suggests that drugs targeting orf3a might be a therapeutic option against a broad range of CoV-related diseases [199].

Recently the SARS-CoV-2 orf3a structure was solved by cryo-EM (PDB 6XDC) [199]. The N-terminus (amino acids 1-39), the C-terminus (239-275) and a short loop (175-180) were not observed, probably due to molecular disorder. Orf3a was solved as a homodimer, although the Authors were able to reconstruct the tetramer at a lower final resolution of ∼6.5 Å, also showing a model of the neighboring dimers [199], and inferring from this model that residues W131, R134, K136, H150, T151, N152, C153, and D155 were involved in a network of interactions that would mediate tetramerization (Figure 11a). Considering that SARS-CoV orf3a has been identified as an emodin-sensitive potassium-permeable cation channel, the narrow size of the pore in the SARS-CoV-2 orf3a structure strongly suggests that the latter is in the closed or inactive conformation [199].

**Figure 11.**
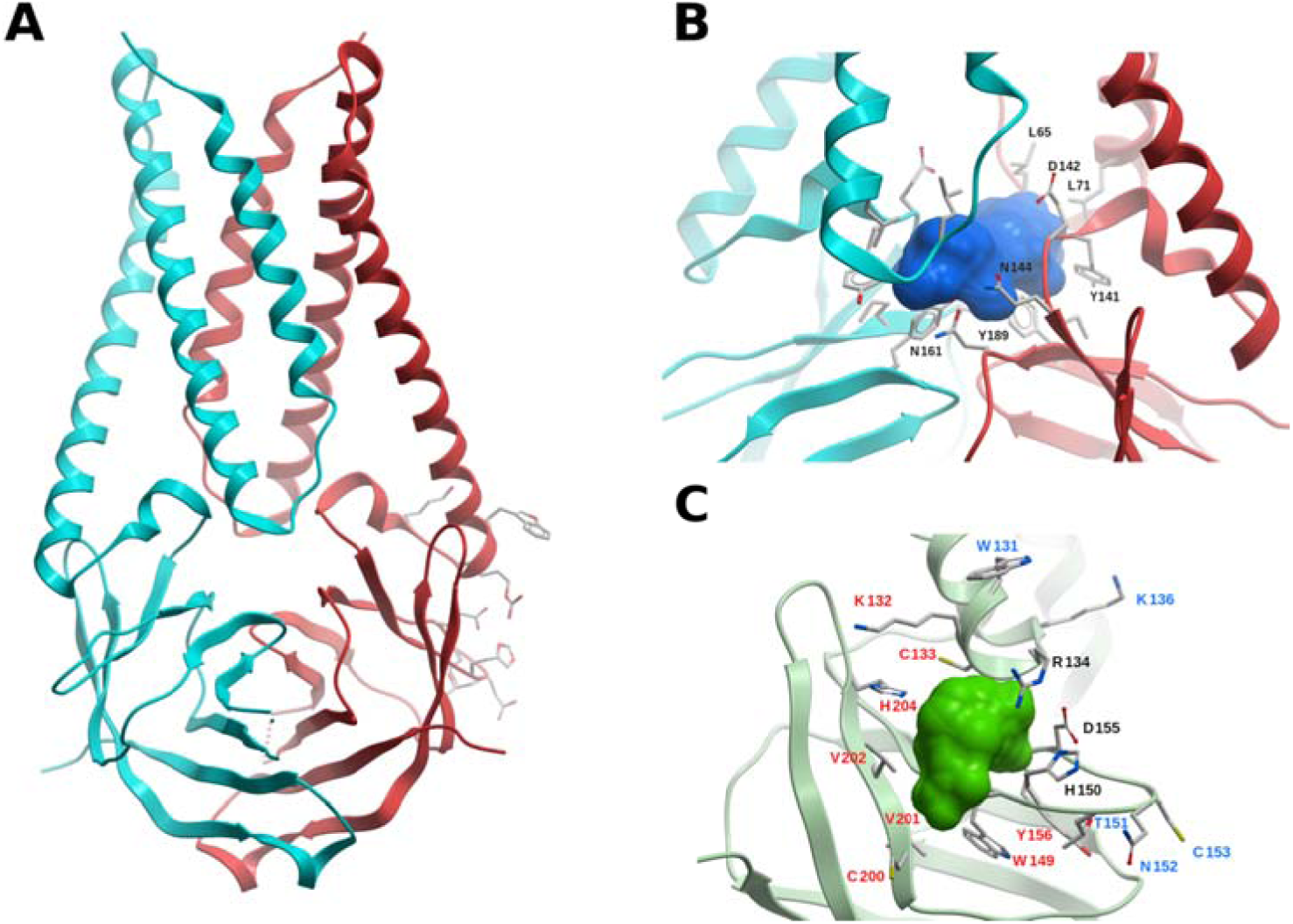
Structure and potential binding sites of the SARS-CoV-2 accessory protein orf3a. (A) Ribbon representation of the orf3a homodimer (cyan and red). Residues W131, R134, K136, H150, T151, N152,C153, and D155, which might be involved in homo-tetramerization, are displayed (not labeled for the sake of clarity). (B) Potential druggable site within the orf3a homodimer interface (blue surface). The neighboring residues are displayed, and labeled for one protomer. (C) Tetramerization interface and a partially overlapping potentially druggable site (green molecular surface). Residue l bels have been colored red (binding site), blue (tetramerization interface), black (common residues).

Using FTMap and ICM Pocket Finder, a druggable site was found within the dimer, lined up by the side chains L65, L71, Y141, D142, N144, P159, N161, and Y189 of both protomers (Figure 11b). This cavity has also been identified in Ref. [199]. Since conformational changes of the TMs are needed for channel opening, we could hypothesize that a small-molecule binding within this site would interfere with this rearrangement, or directly block the channel. Another potentially druggable site delimited by residues K132-R134, C148-H150, D155, Y156, C200-V202, and H204 was identified, which partially overlaps with the proposed tetramerization interface, and thus it could be explored for a possible PPI inhibitor (Figure 11c).

### Orf7a/X4/U122

SARS-CoV-2 orf7a is a transmembrane protein of 121 amino acids (106 if only the mature protein is considered, excluding the signalling peptide), with 86% identity and 94% similarity with respect to its SARS-CoV counterpart. Interestingly, it exhibits structural similarity to Igs [200,201], but shares poor sequence identity with proteins of the Ig superfamily (within the 2%-16% range). Like orf3a, orf7a is also included in the virion, and both proteins have been shown to interact with each other in SARS-CoV [196,202] and in SARS-CoV-2 [166]. In SARS-CoV, orf7a expression increases the expression of NFκB, IL8 and JNK, all involved in inflammatory responses [198], and the deletion of orf7a reduces the virus titer in 30-fold. Orf7a appears to be unique to SARS CoVs, showing no significant similarity to any other protein, either viral or non-viral [201]. In SARS-CoV-2, orf7a interacts with midasin AAA ATPase 1 (MDN1) and HEAT repeat containing 3 (HEATR3) [94]. MDN1 is a protein involved in the maturation of ribosomes in eucariotes [203], while HEATR3 have a positive role in Nucleotide-binding oligomerization domain-containing protein 2 (NOD2) mediated NF-κB signaling [204] and is also involved in the assembly of the 5S ribosomal subunit [205]. From this information it can be expected that orf7a works by regulating inflammation as its SARS-CoV homolog, and also that SARS-CoV-2 orf7 is involved the protein translation regulation. The N-terminal ectodomain structure of orf7a (amino acids 1-66, using the numbering of the mature protein) is available in SARS-CoV (PDB 1XAK at 1.8 Å [201], and PDB 1OY4 solved by NMR [206]), and recently in SARS-CoV-2 (PDB 6W37). The structure displays a seven-stranded β-sandwich fold with two disulfide bonds (Figure S14), and exhibits main chain RMSD values of 0.4 Å and 0.9 Å with the corresponding SARS-CoV orf7a structures, respectively. The SARS-CoV NMR structure also features a disordered part corresponding to the stalk region, between the ectodomain and the membrane (residues 68-82). The transmembrane region comprises approximately amino acids 83-101.

While the functionality and interaction with partners of orf7a are not clearly known, we identified two hot-spots on the surface of orf7a. An extended cryptic site was predicted lined up by residues H4, Q6, L16, P19, Y60, L62, with a CS nearby which includes a deep hydrophobic pocket already identified in SARS-CoV as a potential PPI site [201] (Figure S14). A hot spot defined by residues E18, T24, Y25, F31, P33, A35, N37, and F50 was also identified (Figure S14). Both sites might have functional roles, and further studies are needed to confirm this hypothesis, and whether small-molecules binding to any of them might interfere with the viral mechanism.

### Orf9b

In SARS-CoV, orf9b is a non-essential dimeric membrane protein, produced from an alternative start codon to N-orf9a, with a lipid-binding-like structure [207], and with the ability to bind several other viral proteins [208], including structural proteins, which allows orf9b to be incorporated into the virion [209]. In SARS-CoV, its action is used in interfering with mitochondrial factors that limit IFN responses [210] and in regulating apoptosis, which is a mechanism associated with the immune response [211]. In SARS-CoV-2, it suppresses IFN-I responses through association with TOM70 in mitochondria [212], showing role conservation in SARS-CoV. Additionally, in SARS-CoV-2, orf9b has been reported to interfere with microtubule organization and IRES dependent translation factors [15].

The homodimer structure of SARS-CoV-2 orf9b has been solved by x-ray crystallography at 2.0 Å (PDB 6Z4U); the SARS-CoV structure at 2.8 Å is also available (PDB 2CME). The loop M26-G38 is not present in the structure, likely due to its high flexibility. The sequence identity and similarity of orf9b between SARS-CoV and SARS-CoV-2 is 73% and 83%, respectively, and the main chain RMSD values of the orf9b dimers corresponding to both species (measured using residues with defined secondary structure) is 0.65 Å (a higher value of ∼2 Å is obtained using the full length, due to the high B-factor loops). The structure features a 2-fold symmetric dimer, where both protomers are in a highly interlocked architecture, as in a handshake (Figure S15a).

The dimer exhibits a central hydrophobic cavity (Figure S15a) lined up by residues V15, I19, L21, I44, L46, L52-L54, I74, V76, M78, and V94 of both protomers; in the crystal structure PDB 6Z4U this central cavity is filled with polyethylene glycol (in the corresponding SARS-CoV structure a decane molecule is present). In SARS-CoV, it was hypothesized that orf9b immerses its positively charged surface into the negatively charged lipid head groups of the membrane, while becoming anchored by lipid tails that could bind to this hydrophobic cavity [207]. In this context, a small-molecule which could bind to the central cavity would impede membrane attachment by competing with the lipid tails.

Using FTMap, two potentially druggable sites were identified, where a molecule binding to them might interfere with homodimerization (Figure S15). These sites are defined by residues: i) D2-I5, M8, L12, I45, R47, L87, D89, F91, and V93; ii) V15, P17-L21, V41, I44, L46, S53, L54, V76, and V94; the latter site lies in a region with above-average cryptic site score.

### Other orf accessory proteins

As was described earlier, the polyprotein orf1AB codes for the replicative proteins, DMVs formation, and some processes involved in evading immune innate responses. Orf2, orf4, orf5 and orf9a code for the S, E, M, and N proteins, respectively, which are the four structural virion proteins. SARS-CoV-2 possess at the least other six accessory proteins orf3b,orf6, orf7b, orf8, orf9c/orf14 and orf10, and some of them are produced by alternative start codons of the same orf. These proteins are more divergent among different CoVs than those previously described; since not being essential for replication, they have a lower natural selection pressure, and therefore a higher mutation rate. In fact, the deletion of orf3a, orf3b, orf6, orf7a, and orf7b in SARS-CoV do not abrogate viral production *in vitro* [213], nevertheless it does affect pathogenicity and virulence.

The following proteins lack of experimentally solved structure, or they cannot be modeled due to the absence of similar experimentally solved proteins which could serve as templates:

Orf3b is a short protein limited to some β- and γ-coronaviruses [214], and produced by an alternative start-codon. In SARS-CoV it was shown to inhibit the expression of IFN-β during synthesis and signaling [215]. It differs considerably between different CoVs, but maintains its pathogenic function [216].

The SARS-CoV orf6 is a small membrane protein [217] sharing 69% of identity with its SARS-CoV-2 counterpart. It acts as a pathogenicity factor, due to its capacity of converting a sublethal MHV infection into a lethal one [218]; this property only depends on the N-terminal transmembrane segment [219]. The SARS-CoV orf6 enhances viral replication [220] through interaction with components of the viral replication machinery, such as nsp8 [221]. In addition, it shows pathogenic activity due to its ability to induce apoptosis, similar to orf3a and orf7a [222]. Additionally, SARS-CoV-2 orf6 is involved in the inhibition of IFN signaling [223].

In SARS-CoV, orf7b is a small membrane protein that could be included in the virion [224], and is probably involved in attenuating viral production [225].

Just like orf7a, orf8 is predicted to have an immunoglobulin-like structure [200] from where it can be assumed that orf8 could play roles in immune evasion and pathogenesis. In fact, SARS-CoV-2 orf8 is involved in inhibiting IFN signaling [223]. Moreover, orf8 is the fastest evolving gene in SARS-CoV-2, as can be inferred by its high variability [200].

Orf9c/orf14 is the third putative protein translated from the orf9 [226], and the least characterized one. There is only proteomic data, that shows its interaction with factors associated with mitochondrial regulation, cytokine production, and coagulation [15].

Orf10 is a putative membrane protein present only in SARS related CoVs [227]. No functional activity has been determined experimentally, but it is inferred from proteomic data that it is involved in targeting host proteins for proteasome degradation [15].

## 4 Discussion and Perspective

The recent appearance of COVID-19 caused by the SARS-CoV-2, its fast spread throughout the world, and the mounting number of infected persons have triggered a prompt and resolute quest for therapeutic options to treat this serious infectious disease.

A vaccine capable of generating a protective immune response would be the best option to control this disease, but the above stated factors, coupled with the risk of death and the threat to the national health systems –especially in low-income countries– highlight the necessity of the quick development of a treatment to cure this disease, or at least to control its severity.

Although there are no specific antivirals available, approved or experimental drugs are being evaluated through drug repurposing strategies [18-23], mainly targeting the two viral cysteine proteases and the polymerase complex. However, protease inhibitors might lack selectivity [24], and the efficacy of nucleoside inhibitors targeting the RdRp is limited by the ExoN proofreading machine and non-mutagenic doses limitations, and thus alternative or complementary therapeutic strategies to fight COVID-19 are needed.

In this work we stress the fact that besides the replicative and structural proteins, all of which are critical for the viral cycle, other proteins also have vital functions, and thus would constitute excellent targets for drug discovery. The helicase (nsp13), and methyl-transferases N7-Mtase (nsp14) and 2′-O-MTase (nsp16) contribute to genome stability through their involvement in the capping process; the ExoN (nsp14) is responsible for proofreading, and thus for the extremely low mutation rate and nucleoside analogs resistanse of SARS-CoV-2; several nsp3 domains, such as SUD, NAB and Ub1 are known to bind ssRNA; the NendoU (nsp15) cleaves polyuridines produced during the priming of the poly(A) ssRNA during replication, which helps to dampen dsRNA MDA5-dependent antiviral IFN responses [125]; nsp9 acts as a hub that binds ssRNA and interacts with nsp8, the N protein, and several host nuclear pore proteins [15,130,131]; the ADRP and PL^pro^ attenuate the effects of IFN and cytokine signaling components that induce antiviral and inflammatory responses [54,142]; orf3a and orf7a induce NFκB, IL-8, and JNK, promoting inflammatory responses [198], while orf3a also induces the production of fibrinogen, promoting fibrosis, one of the complications of COVID-19 [197]; orf9b is involved in suppressing mitochondrial mediated IFN antiviral responses [210]; orf6 is known to increase the lethality in CoVs by enhancing viral replication and inhibiting IFN signaling [220,223]. In CoVs, a delayed IFN response is a redundant pattern that allows robust viral replication, and also induces the accumulation of cytokine-producing macrophages, thus increasing the severity of the disease [228].

Given this scenario, a thorough characterization of the druggability of the SARS-CoV-2 proteins provides an array of alternative targets for drug discovery. We present an in-depth functional, structural and druggability analysis of all non-structural, structural, and accessory proteins of SARS-CoV-2, identifying potential druggable allosteric and PPI sites throughout the whole proteome, thus broadening the repertoire of current targetable proteins. It should be stressed that druggability characterization of a site does not necessarily imply that any compound binding at that site will modulate that target and exhibit an observable biological effect.

We are convinced that our work will contribute to the quick development of an effective SARS-CoV-2 antiviral strategy, which in view of the high similarity among CoVs, it might be useful to fight related viruses. Moreover, these therapeutic options might be instrumental in fighting different CoV-associated diseases that could threaten global health in the future.

## 5 Materials and Methods

### 5.1 Molecular system setup

All structures were downloaded from the Protein Data Bank (PDB) and prepared using the ICM software [229] (MolSoft LLC, San Diego, CA, 2019) in a similar fashion as in earlier works [230]. Succinctly, hydrogen atoms were added, followed by a short local energy minimization in the torsional space; the positions of polar and water hydrogens were determined by optimizing the hydrogen bonding network, and then all water molecules were deleted. All Asp and Glu residues were assigned a -1 charge, and all Arg and Lys residues were assigned a +1 charge. Histidine tautomers were chosen according to their corresponding hydrogen bonding pattern.

### 5.2 Homology modeling

In each case, a crude model was built using the backbone structure of the template, and then refined through local energy minimization using ICM. To avoid pocket collapse, and taking into account the complete binding site conservation, whenever available, ligands were kept within the binding site during the refinement process, in a ligand-steered modeling fashion [231-233].

### 5.3 Hot spots and cryptic sites

Identification of binding energy hot spots was performed with FTMap (https://ftmap.bu.edu) [26,27]. The method samples through rigid docking a library of 16 small organic probe molecules of different size, shape and polarity on the protein. For each probe, all the poses generated are clustered using a 4 Å clustering radius, and then clusters are ranked on the basis of their average energy, keeping the six lowest-energy clusters for each probe. After the probe clusters of all the 16 molecules have been generated, they are then re-clustered based on vicinity into consensus sites (CSs). These CSs (hot spots) are ranked on the basis of the number of their probe clusters. The program offers a protein-protein interaction (PPI) mode, where the aim is to identify hot spots on protein-protein interfaces. To identify binding sites, the top ranking CS is considered the kernel of the binding site, and is expanded by adding neighboring CSs with a center-to-center distance (CD) of less than 8 Å to any existing CS in the binding site, until no further expansion is possible. The binding site is defined as those residues within 4 Å of the probes of the CSs used to describe the binding site. The top ranking CS is removed and the procedure repeated starting from the second ranking CS, and so forth. Considering the druggability criteria (see below), only CSs with at least 13 probe clusters were considered to be expanded.

Cryptic sites on proteins were determined using CryptoSite [30] (https://modbase.compbio.ucsf.edu/cryptosite/); cryptic sites are those formed only in ligand-bound structures, but usually “hidden” in unbound structures. Taking into account the analysis of the druggability of the cryptic sites [234], only those cryptic sites with at least 16 probe clusters (determined with FTMap) within 5 Å were considered as potentially druggable [235].

The ICM Pocket Finder method predicts the position and shapes of cavities and clefts using a transformation of the Lennard-Jones potential by convolution with a Gaussian kernel of a certain size, and construction of equipotential surfaces along the maps of a binding potential [29,236].

### 5.4 Druggability criteria

The druggability of a site was characterized based on the the CSs generated by FTMap in terms of: i) the number of probe clusters in the primary hot spot (S), ii) if there are one or more secondary spots with a CD < 8 Å from the primary spot, and iii) the maximum dimension (MaxD) of the connected ensemble (measured as the distance between the two most separated probe atoms within the probe clusters) [31]. In general, a site was considered druggable if S ≥ 16, CD < 8 Å, and MaxD ≥ 10 Å; non-druggable if S < 13 or MaxD < 7 Å; borderline druggable if 13 ≤ S < 16 and CD < 8 Å, or 13 ≤ S < 16, CD ≥ 8 Å, and MaxD ≥ 10 Å.

## Acknowledgments

The Authors thank M. Laura Fernández, Pilar Cossio, and Lucia Cavasotto for critical reading of the manuscript. This work was supported by the National Agency for the Promotion of Science and Technology (ANPCyT) (PICT-2017-3767). CNC thanks Molsoft LLC (San Diego, CA) for providing an academic license for the ICM program.

## Author contributions

C.N.C. conceived the original idea of the work, designed the research process, performed the functional, structural and druggability analyses, interpreted the results and wrote the manuscript. M.S.L. contributed to the functional analysis, interpretation of results, and to the manuscript. J.M. contributed to the analysis, interpretation of results, and to the manuscript. All authors have reviewed the manuscript and approved the submission.

