## Supplementary Information for "Functional and druggability analysis of the SARS-CoV-2 proteome"

**Table of Contents**

Tables S1-S5............................................. 2-8
Figure S1-S15......................................... 9-17
References.................................................18

**Table S1: Domain distribution and available structures of the SARS-CoV-2 nsp3**

| Domain | Aminoacid positions (aproximate) | PDB ID | Identity (%), similarity (%)[b] | Reference |
|---|---|---|---|---|
| Ubl1[a] | 1-112 | 2GRI | 79%, 93% | [1] |
| DUF3655/USD | 113-183 | - | | |
| ADRP/Mac1 | 205-379 | 6VXS, 6WEN, 6WCF, 6W6Y, 6W02<br>6WOJ | | [2]<br>[3] |
| SUD-N/Mac2[a] | 413-540 | 2W2G | 71%, 91% | [4] |
| SUD-M/Mac3[a] | 551-676 | 2W2G<br>2KQV<br>2JZD,2JZF, 2JZE, 2RNK | 79%, 94% | [4]<br>[5]<br>[6] |
| SUD-C/DPUP [a] | 678-746 | 2KQW, 2KAF | 72%, 92% | [5] |
| Ubl2-PL$^{pro}$ | 744-1061 | 6WUU, 6WX4<br>6XA9, 6XAA<br>6W9C, 6WZU, 6WRH,6XG3<br>7JN2, 7JIT, 7JIR, 7JIV, 7JIW | | [7]<br>[8] |
| NAB[a] | 1084-1201 | 2K87 | 81%, 94% | [9] |
| TM-lumen domain-TM/Ectodomain | 1494-1563 | - | | |
| C-Terminal domain/Y-Domain | 1564-1945 | - | | |

[a]No SARS-CoV-2 structure available, the corresponding to SARS-CoV reported.

[b]Relative to the corresponding SARS-CoV-2 nsp3 domain

**Table S2. Available crystallographic structures of the SARS-CoV-2 main protease (M$^{pro}$, nsp5)** (the +100 structures with fragment-like molecules bound to M$^{pro}$ are not included in this table, cf. PDBs 5RGK, 5RF1, 5RG3, and others [10]).

| PDB ID | Ligand PDB ID | Ligand name | Resolution (Å) | Reference |
|---|---|---|---|---|
| 6ZRU | u5g | Boceprevir (bound form) | 2.10 | |
| 6ZRT | sv6 | Telaprevir | 2.10 | |
| 7C7P | sv6 | Telaprevir | 1.74 | |
| 7JFQ | | | 1.55 | |
| 6XQT | nna | (1R,2S,5S)-3-[N-({1-[(tert-butylsulfonyl)methyl]cyclohexyl}carbamoyl)-3-methyl-L-valyl]-N-{(1S)-1-[(1R)-2-(cyclopropyla mino)-1-hydroxy-2-oxoethyl]pentyl}-6,6-dimethyl-3-azabicyclo[3.1.0]hexane-2-carboxamide | 2.30 | |
| 6XQS | sv6 | Telaprevir | 1.90 | |
| 6XQU | u5g | Boceprevir (bound form) | 2.20 | |
| 6XOA | | | 2.10 | |
| 6XMK | qys | (1S,2S)-2-[(N-{[(4,4-difluorocyclohexyl)methoxy]carbonyl}-L-leucyl)amino]-1-hydroxy-3-[(3S)-2-oxopyrrolidin-3-yl]propane-1-sulfonic acid | 1.70 | |
| 6XHM | v2m | N-[(2S)-1-({(2S)-4-hydroxy-3-oxo-1-[(3S)-2-oxopyrrolidin-3-yl]butan-2-yl}amino)-4-methyl-1-oxopentan-2-yl]-4-methoxy-1H-indole-2-carboxamide | 1.41 | |
| 6XKF | | | 1.80 | |
| 6XKH | | | 1.28 | |
| 6XHU | | | 1.80 | |
| 6XFN | UAW243 | UAW243 | 1.70 | |
| 7C8U | k36 | GC376 | 2.35 | |
| 6Z2E | q5t | (4~{S})-4-[[(2~{S})-2-[[(2~{S})-2-[[(2~{S})-2-[3-[2-[2-[2-[2-[5-[(3~{a}~{S},4~{R},6~{a}~{R})-2-oxidanylidene-3,3~{a},4,6~{a}-tetrahydro-1~{H}-thieno[3,4-d]imidazol-4-yl]pentanoylamino]ethoxy]ethoxy]ethoxy]ethoxy]propanoylamino]butanoyl]amino]-3,3-dimethyl-butanoyl]amino]-4-methyl-pentanoyl]amino]-6-methylsulfonyl-hexanamide | 1.70 | |
| 6XA4 | UAW241 | UAW241 | 1.65 | |
| 6XB1 | nen | 1-ethyl-pyrrodoline-2,5-dione | 1.80 | |
| 6XB0 | | | 1.80 | |

| PDB ID | Ligand PDB ID | Ligand name | Resolution (Å) | Reference |
|---|---|---|---|---|
| 6XB2 | nen | 1-ethyl-pyrrodoline-2,5-dione | 2.10 | |
| 6XBI | UAW248 | UAW248 | 1.70 | |
| 6XBH | UAW247 | UAW247 | 1.60 | |
| 6XBG | UAW246 | UAW246 | 1.45 | |
| 6XCH | ar7 | Leupeptin | 2.20 | |
| 7C8T | nol | N-[(BENZYLOXY)CARBONYL]-O-(TERT-BUTYL)-L-THREONYL-3-CYCLOHEXYL-N-[(1S)-2-HYDROXY-1-{[(3S)-2-OXOPYRROLIDIN-3-YL]METHYL}ETHYL]-L-ALANINAMIDE | 2.05 | |
| 7C8R | tg3 | ethyl (4R)-4-[[(2S)-4-methyl-2-[[(2S,3R)-3-[(2-methylpropan-2-yl)oxy]-2-(phenylmethoxycarbonylamino)butanoyl]amino]pentanoyl]amino]-5-[(3S)-2-oxidanylidenepyrrolidin-3-yl]pentanoate | 2.30 | |
| 6YVF | | | 1.60 | |
| 6WTK | ued | N~2~-[(benzyloxy)carbonyl]-N-{(2S)-1-hydroxy-3-[(3S)-2-oxopyrrolidin-3-yl]propan-2-yl}-L-leucinamide | 2.00 | |
| 6WTM | | | 1.85 | |
| 6WTJ | k36 | GC376 | 1.90 | |
| 6YZ6 | PRD_000216 | Leupeptin | 1.70 | |
| 6WTT | k36 | GC376 | 2.15 | |
| 7BRR | k36 | GC376 | 1.40 | |
| 7BRO | | | 2.00 | |
| 7BRP | hu5 | Boceprevir | 1.80 | |
| 6WNP | u5g | Boceprevir (bound form) | 1.44 | |
| 6YT8 | pk8 | Zinc pyrithione | 2.05 | |
| 6WQF | | | 2.30 | |
| 6M0K | fjc | ~{N}-[(2~{S})-3-(3-fluorophenyl)-1-oxidanylidene-1-[[(2~{S})-1-oxidanylidene-3-[(3~{S})-2-oxidanylidenepyrrolidin-3-yl]propan-2-yl]amino]propan-2-yl]-1~{H}-indole-2-carboxamide | 1.50 | [11] |
| 6LZE | fhr | ~{N}-[(2~{S})-3-cyclohexyl-1-oxidanylidene-1-[[(2~{S})-1-oxidanylidene-3-[(3~{S})-2-oxidanylidenepyrrolidin-3-yl]propan-2-yl]amino]propan-2-yl]-1~{H}-indole-2-carboxamide | 1.51 | [11] |
| 7BUY | jry | Carmofur | 1.60 | [12] |
| 6YNQ | p6n | 2-methyl-1-tetralone | 1.80 | |

| PDB ID | Ligand PDB ID | Ligand name | Resolution (Å) | Reference |
|---|---|---|---|---|
| 7BQY | PRD_002214 | N3 | 1.70 | [13] |
| 6M2N | 3wl | 5,6,7-trihydroxy-2-phenyl-4H-chromen-4-one | 2.20 | |
| 6M2Q | | | 1.70 | |
| 6W63 | x77 | N-(4-tert-butylphenyl)-N-[(1R)-2-(cyclohexylamino)-2-oxo-1-(pyridin-3-yl)ethyl]-1H-imidazole-4-carboxamide | 2.10 | |
| 6YB7 | | | 1.27 | |
| 6M03 | | | 2.00 | |
| 6Y84 | | | 1.39 | |
| 6Y2G | o2k | α-ketoamide | 2.20 | [14] |
| 6Y2F | o2k | α -ketoamide | 1.95 | [14] |
| 6Y2E | | | 1.75 | [14] |
| 6LU7 | PRD_002214 | N3 | 2.16 | [13] |

**Table S3. Available experimental structures of the SARS-CoV-2 RdRp complex (nsp12-nsp8-nsp7)**

| PDB ID | PDB Structure Title | Resolution (Å)[a] | Reference |
|---|---|---|---|
| 6XQB | SARS-CoV-2 RdRp/RNA complex | 3.40 | |
| 6XEZ | Structure of SARS-CoV-2 replication-transcription complex bound to nsp13 helicase - nsp13(2)-RTC | 3.50 | [15] |
| 7BV1 | Cryo-EM structure of the apo nsp12-nsp7-nsp8 complex | 2.80 | [16] |
| 6X2G | SARS-CoV-2 RdRp/RNA complex | 3.53 | |
| 7BV2 | The nsp12-nsp7-nsp8 complex bound to the template-primer RNA and triphosphate form of Remdesivir(RTP) | 2.50 | [16] |
| 6YYT | Structure of replicating SARS-CoV-2 polymerase | 2.90 | [17] |
| 7BZF | COVID-19 RNA-dependent RNA polymerase post-translocated catalytic complex | 3.26 | [18] |
| 7C2K | COVID-19 RNA-dependent RNA polymerase pre-translocated catalytic complex | 2.93 | [18] |
| 6M71 | SARS-Cov-2 RNA-dependent RNA polymerase in complex with cofactors | 2.90 | [19] |
| 7BW4 | Structure of the RNA-dependent RNA polymerase from SARS-CoV-2 | 3.70 | [20] |
| 7BTF | SARS-CoV-2 RNA-dependent RNA polymerase in complex with cofactors in reduced condition | 2.95 | [19] |

[a]All structures have been solved by cryo-EM

**Table S4. Available experimental structures of the RNA nucleoside-2'O-methyltransferase complex (nsp16-nsp10)**

| PDB ID | Ligand PDB ID | Ligand name | Resolution[a] (Å) | Reference |
|---|---|---|---|---|
| 6XKM | sam | SAM[b] | 2.25 | |
| 7BQ7 | sam | SAM | 2.37 | |
| 6WKS | sam<br>gta | SAM<br>P1-7-methyguanosine-P3-adenosine-5’,5’-triphosphate | 1.80 | [21] |
| 6YZ1 | sfg | Sinefungin | 2.40 | [22] |
| 6W4H | sam | SAM | 1.80 | [23] |
| 6W75 | sam | SAM | 1.95 | [23] |
| 6W61 | sam | SAM | 2.00 | [23] |
| 6WKQ | sfg | Sinefungin | 1.98 | |
| 6WVN | sam<br>gta | SAM<br>P1-7-methyguanosine-P3-adenosine-5’,5’-triphosphate | 2.00 | |
| 7C2I | sam | SAM | 2.50 | |
| 7C2J | sam | SAM | 2.80 | |
| 6WJT | sah | SAH[c] | 2.00 | |
| 6WQ3 | sah<br>gta<br>8nk | SAH<br>P1-7-methyguanosine-P3-adenosine-5’,5’-triphosphate<br>7-methylguanosine 5'-diphosphate | 2.10 | |
| 6WRZ | sah<br>gta<br>mgp | SAH<br>P1-7-methyguanosine-P3-adenosine-5’,5’-triphosphate<br>7-methyguanosine- 5’-triphosphate | 2.25 | |

[a]All structures have been solved by crystallography
[b]S-adenosyl-L-methionine
[c]S-adenosyl-L-homocysteine

**Table S5. Available experimental structures of the SARS-CoV-2 nucleocapsid N protein**

| PDB ID | N Protein domain | Resolution (Å)[a] | Reference |
|---|---|---|---|
| 6WZQ | Dimerization (C-terminal) | 1.45 | [24] |
| 6WZO | Dimerization (C-terminal) | 1.42 | [24] |
| 6YI3 | RNA-binding domain (N-terminal) | NMR | |
| 7C22 | Dimerization (C-terminal) | 2.00 | |
| 6VYO | RNA-binding domain (N-terminal) | 1.70 | |
| 6M3M | RNA-binding domain (N-terminal) | 2.70 | [25] |
| 6WKP | RNA-binding domain (N-terminal) | 2.67 | |
| 6YUN | Dimerization (C-terminal) | 1.44 | |
| 6ZCO | Dimerization (C-terminal) | 1.36 | |
| 6WJI | Dimerization (C-terminal) | 2.05 | |

[a]All structures have been solved by crystallography except were noted.

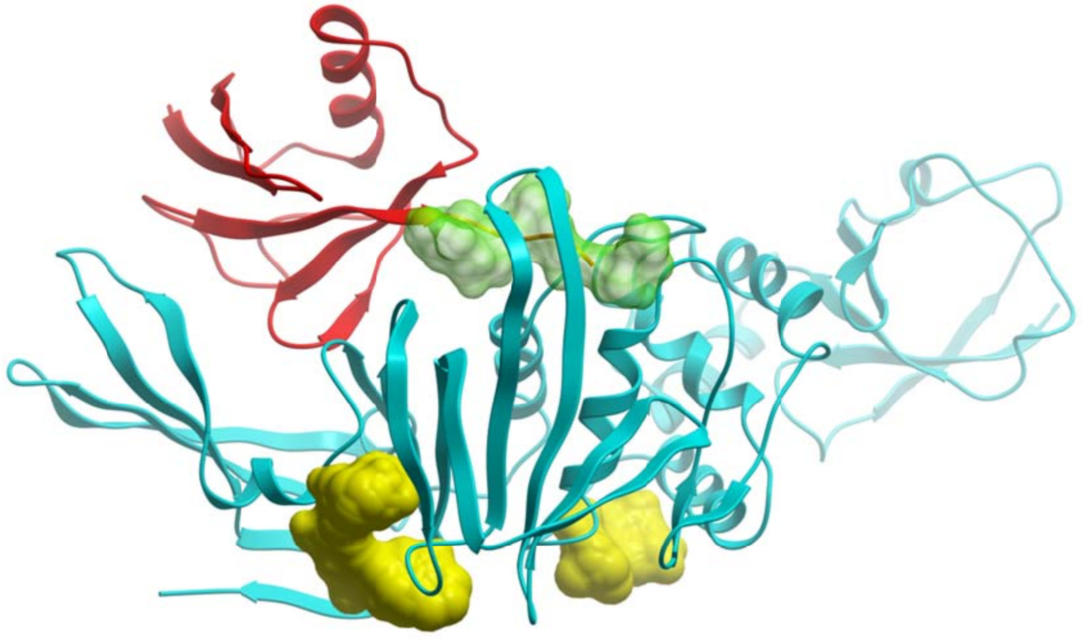

**Figure S1. Structure of the SARS-CoV-2 PL$^{pro}$.** Overall structure of PL$^{pro}$ (cyan ribbon), in complex with ubiquitin (red ribbon). The N-terminal part of ubiquitin fits within the catalytic binding site (transparent surface). Two potential binding sites are displayed in yellow.

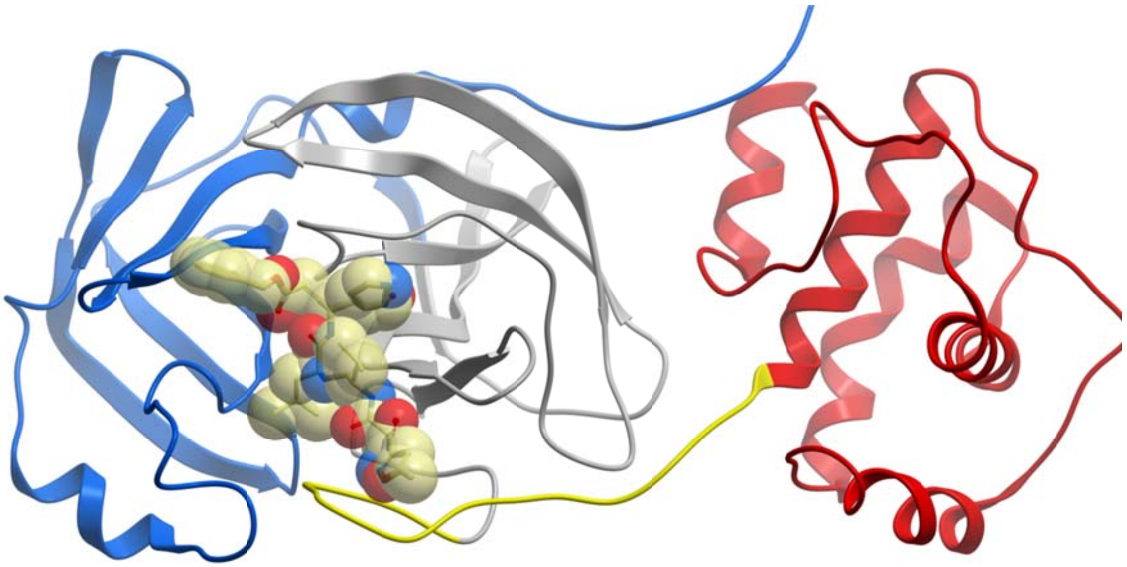

**Figure S2. Structure of SARS-CoV-2 M$^{pro}$.** Overall structure of M$^{pro}$ (cyan ribbon), in complex with peptide inhibitor N3 (light yellow carbon atoms). Color code: Domain I, blue; Domain II, gray; Domain III, red; Domains II-III loop, yellow.

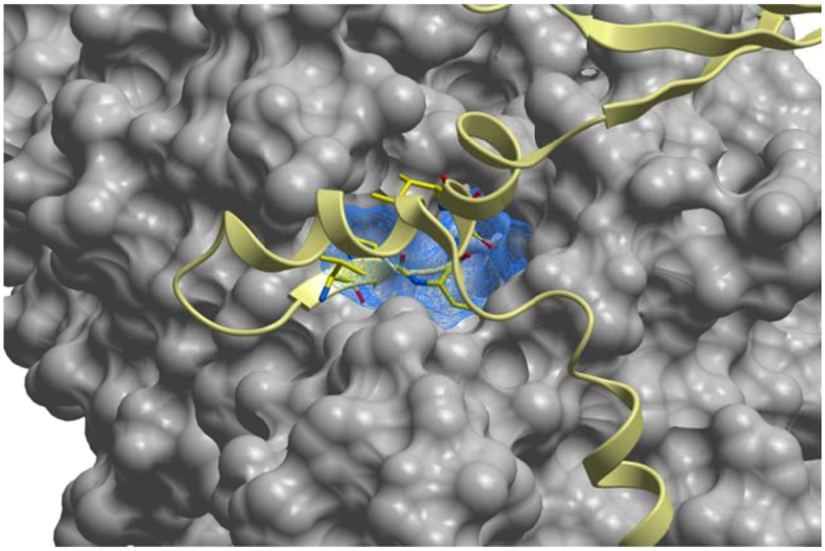

**Figure S3: Predicted potential druggable site in nsp12.** Target site identified using FTMap in nsp12 (grey surface). The predicted binding site is displayed in blue mesh representation, and nsp8 is shown as yellow ribbon. Nsp8 residues V115-I119 are displayed (though not labeled as sticks) to highlight that a small molecule binding to these potentials sites might interfere with PPI.

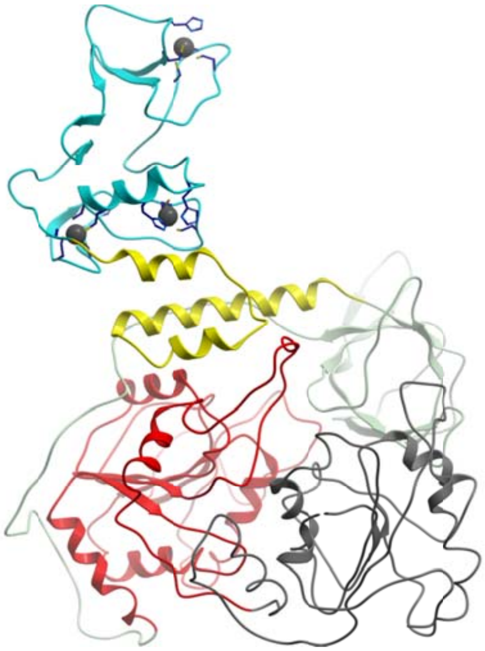

**Figure S4: Overall structure of the SARS-CoV-2 helicase/NTPase/Rtpase (nsp13).** Domain color code: zinc binding domain, cyan; stalk, yellow; 1B, green; 1A, red; 2A, grey. Three zinc atoms are displayed as dark grey spheres. The residues forming the zinc fingers are displayed.

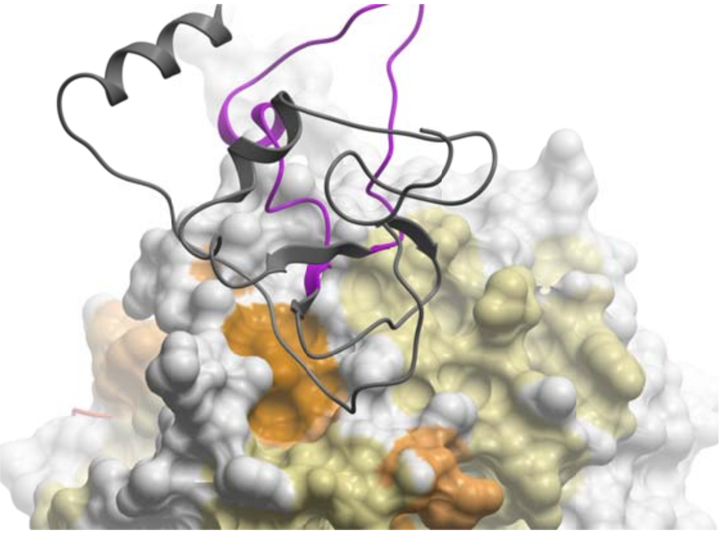

**Figure S5. Potential sites on the SARS-CoV-2 nsp16 surface.** Cryptic site on the PPI surface of nsp16 with nsp10. Nsp16 is colored according to its cryptic site score (score color code: white, low; light brown, intermediate; darker brown, high). Nsp10 is displayed in grey ribbon, with the residues of the 29-mer peptide known to inhibit nsp16-nsp10 interaction colored in magenta.

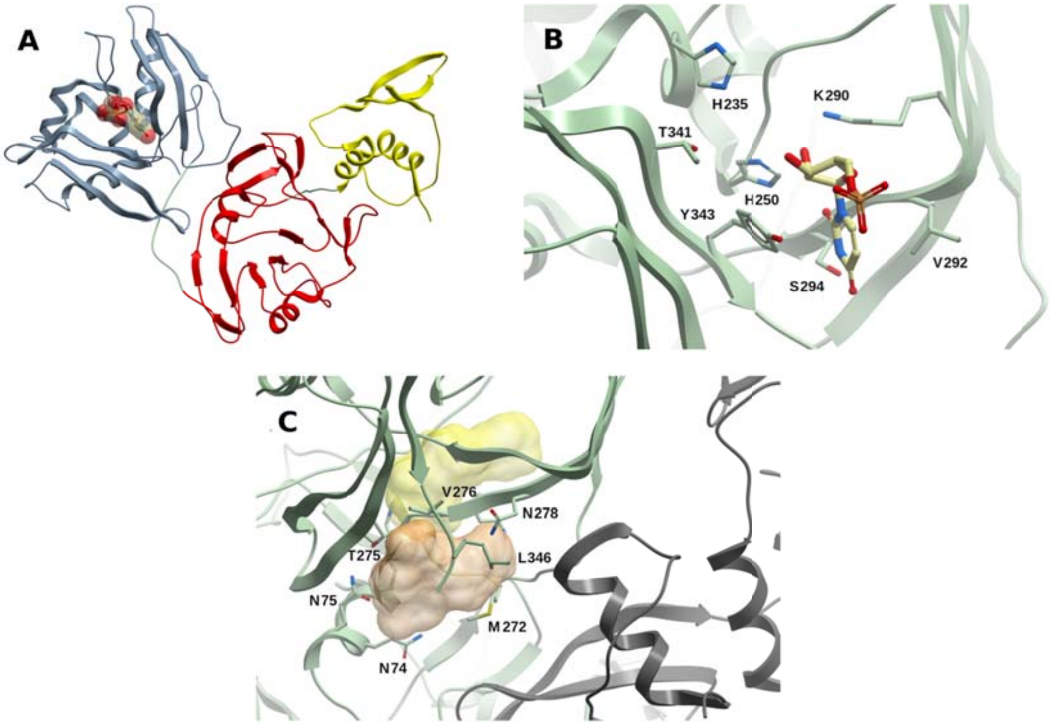


**Figure S6: Structure and binding sites of the SARS-CoV-2 RNA uridylate-specific endoribonuclease (NendoU, nsp15)** (A) Overall structure of nsp15, exhibiting the N-terminal domain (yellow), the central domain (red), and the C-terminal (catalytic) domain (light blue). As reference, a uridine-5'-monophosphate molecule has been placed within the catalytic site. (B) Catalytic site of nsp15 in complex with a uridine-5'-monophosphate molecule (PDB 6WLC). (C) Potential binding sites determined with FTMap colored in yellow (site 1) and tan (site 2). Site 1 is buried, with an opening towards the hexamer channel. Site 2 is delimited by residues of the C-terminus, and closer to another protomer within the nsp15 hexamer (gray ribbon).

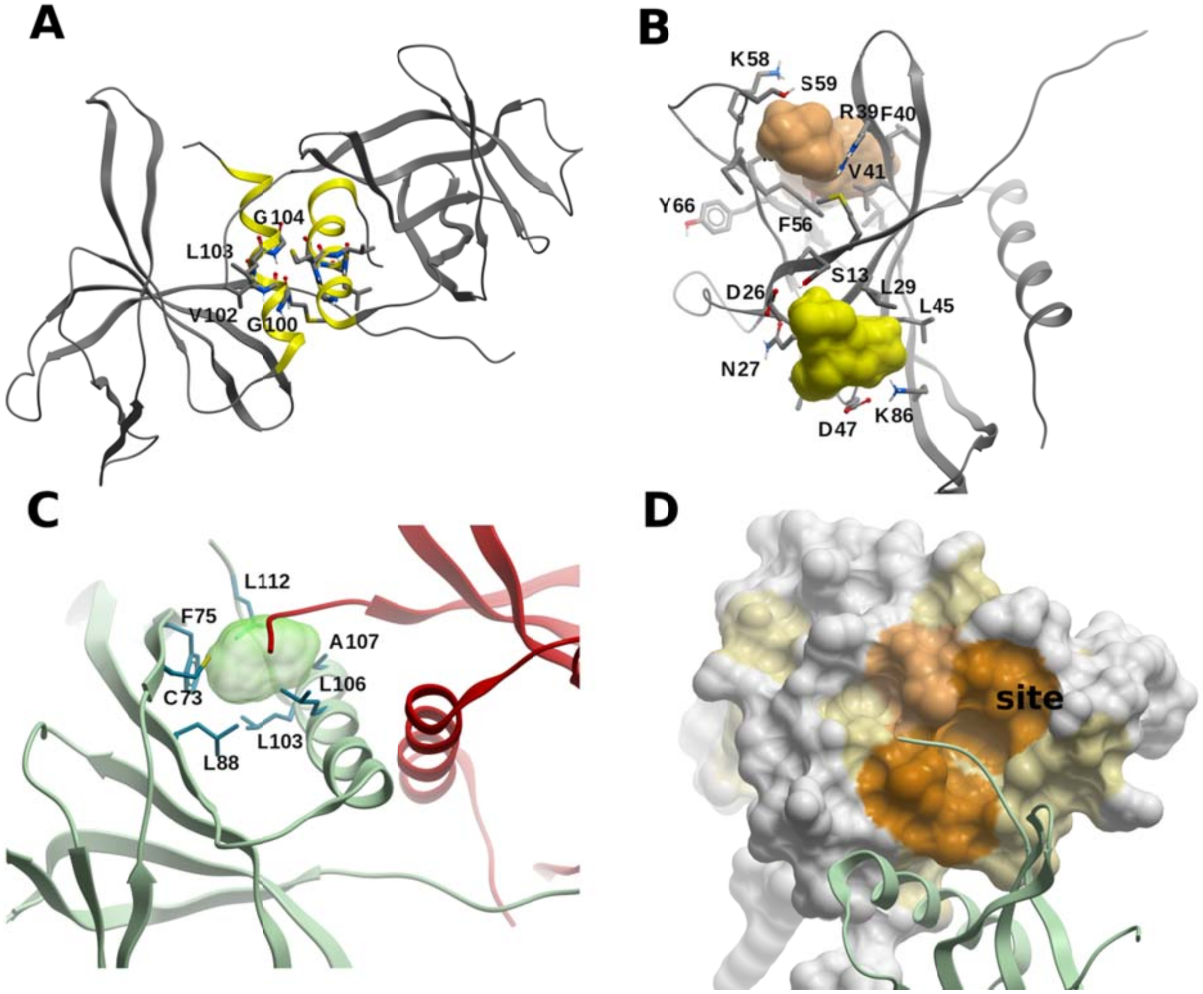


**Figure S7: Structure and potential sites within the SARS-CoV-2 nsp9 homodimer.** (A) Homodimer of nsp9 (PDB 6WXD). The α-helices on each protomer are displayed in yellow, and the residues of the motif GXXXG within those helices are displayed and labeled on one of the monomers (M101 is not labeled for the sake of clarity). (B) Potential druggable sites in nsp9 identified using FTMap, shown in yellow and green. (C) Consensus site within the N-terminus of nsp9 (green ribbon); a small-molecule in this site might interfere with the binding of the other protomer (shown as red ribbon). (D) A different view of the site of (C), showing a high cryptic site score area (color code: white, low; light brown, intermediate; darker brown, high); the other protomer shown as green ribbon.

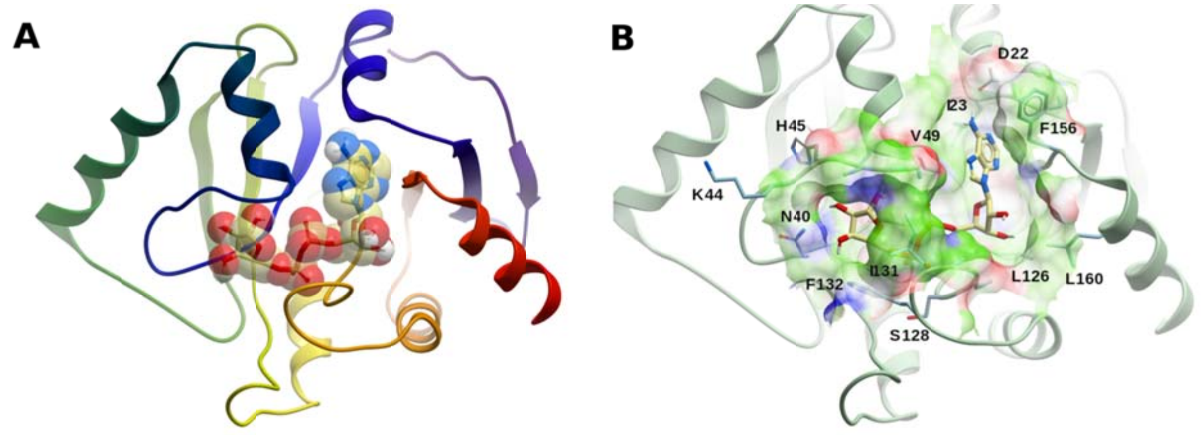


**Fig S8: Structure and binding site of the SARS-CoV-2 ADRP (nsp3 domain).** (A) Overall structure of ADRP in complex with ADP-ribose, colored from the N-terminus (blue) to the C-terminus (red) (PDB 6W02); (B) Binding site of ADRP displaying ADP-ribose; the surface is colored as: green, hydrophobic; red, hydrogen bond acceptor; blue, hydrogen bond donor. Selected residues are labeled.

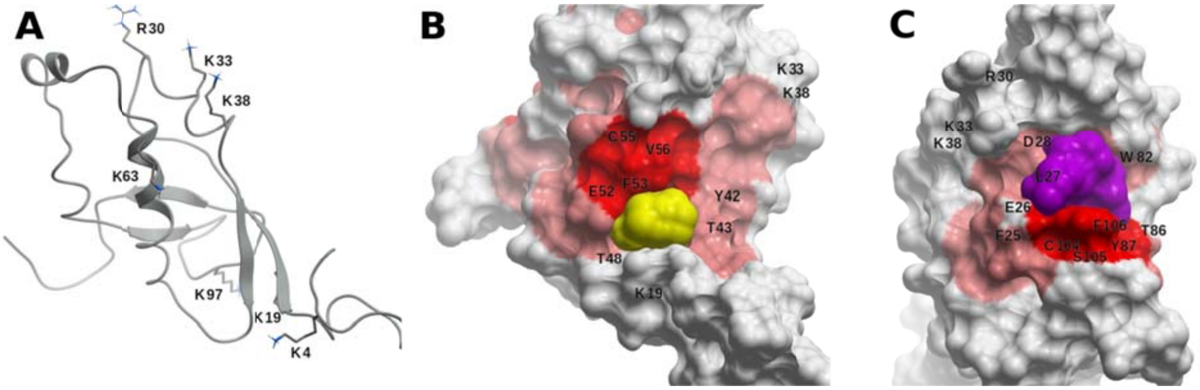


**Figure S9: Structural model of the SARS-CoV-2 Ubiquitin-like 1 nsp3 domain (Ubl1).** (A) Homology model of the nsp3 sub-domain Ubl1 built using the SARS-CoV Ubl1 NMR structure (PDB 2GRI) as template. Positive charged residues which may be involved in RNA binding are displayed and labeled. (B,C) Small potential druggable sites (B, yellow; C, cyan) within areas of above-average cryptic site score (score color code: white, low; light red, intermediate; red, high). Neighboring amino acids are labeled.

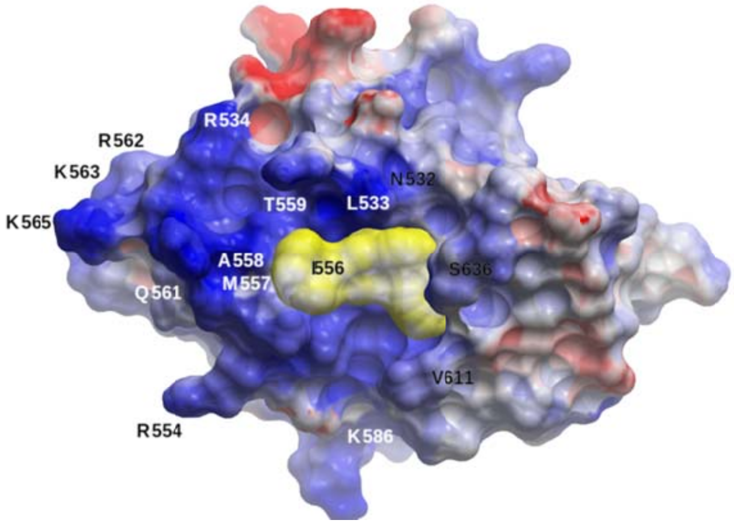


**Figure S10: Structural model of the SARS-CoV-2 SUD-M nsp3 domain.** Electrostatic potential surface of the homology model of SUD-M. The labeled residues define the RNA-binding molecular surface, conserved from SARS-CoV (positive and negative electrostatic areas are represented in blue and red, respectively, neutral in white). A potential druggable binding site is shown in yellow.

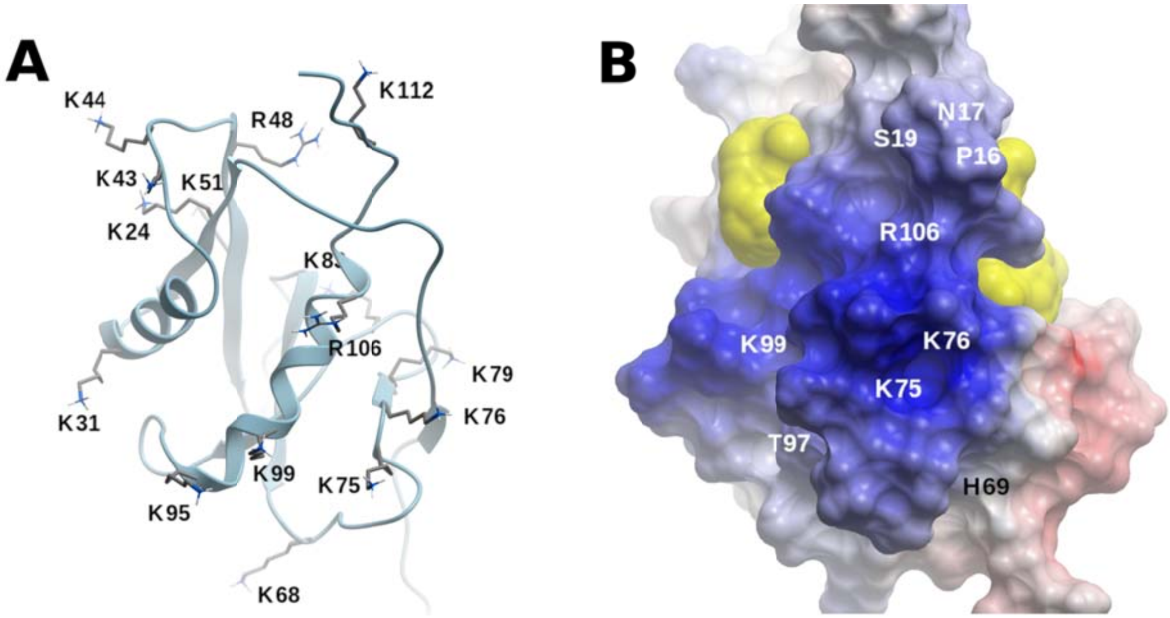


**Figure S11: Structural model of the SARS-CoV-2 nucleic acid binding (NAB) domain of nsp3.** (A) Homology model of the SARS-CoV-2 nsp3 sub-domain NAB built using the SARS-CoV NAB NMR structure (PDB 2K87). Positively charged residues are displayed and labeled. (B) Representation of the electrostatic potential surface of NAB, labeling selected residues identified in ssRNA binding to SARS-CoV NAB (positive and negative electrostatic charges are represented in blue and red, respectively). Two potentially druggable sites, at the sides of the positive charged patch, are shown as yellow surface.

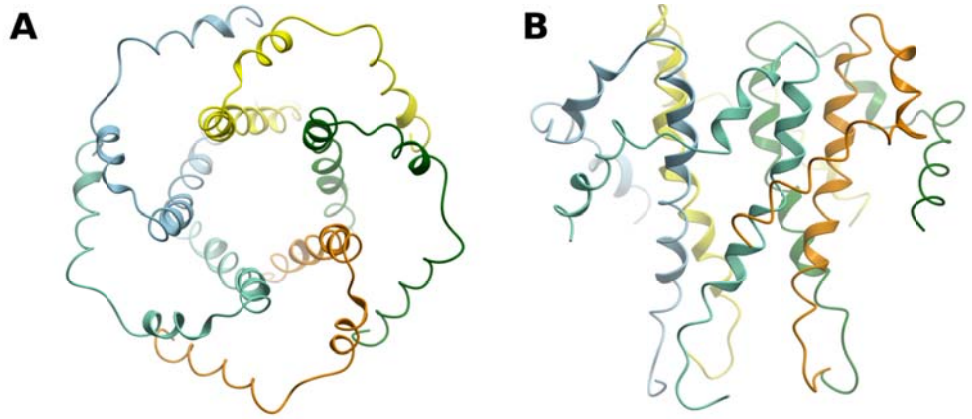


**Figure S12. Structural model of the SARS-CoV-2 E protein homopentamer.** (A) Top-view, and (B) side-view of the structure. Each protomer has been assigned a different color.

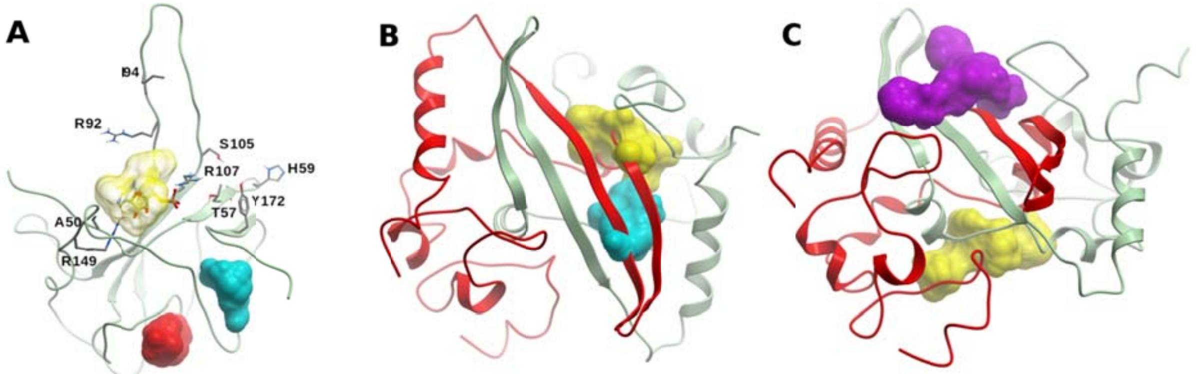


**Figure S13. Structure and potential binding sites of the NTD and CTD of the SARS-CoV-2 N protein.** (A) NTD ribbon representation displaying residues involved in RNA binding. A druggable site is shown in transparent yellow. An AMP molecule from the HcoV-OC43 NTD is also shown. Two potential allosteric sites (red and cyan surfaces) were also identified. (B) CTD dimer (protomer 1, green; protomer 2, red). Two druggable sites on protomer 1 which may interfere with dimer formation are displayed (yellow and cyan molecular surface). (C) Two potential sites predicted on the surface of the CTD dimer (same coloring as in B). One site lies over the central four-stranded β-sheet, and the other opposite to it, near the C-terminal α-helices.

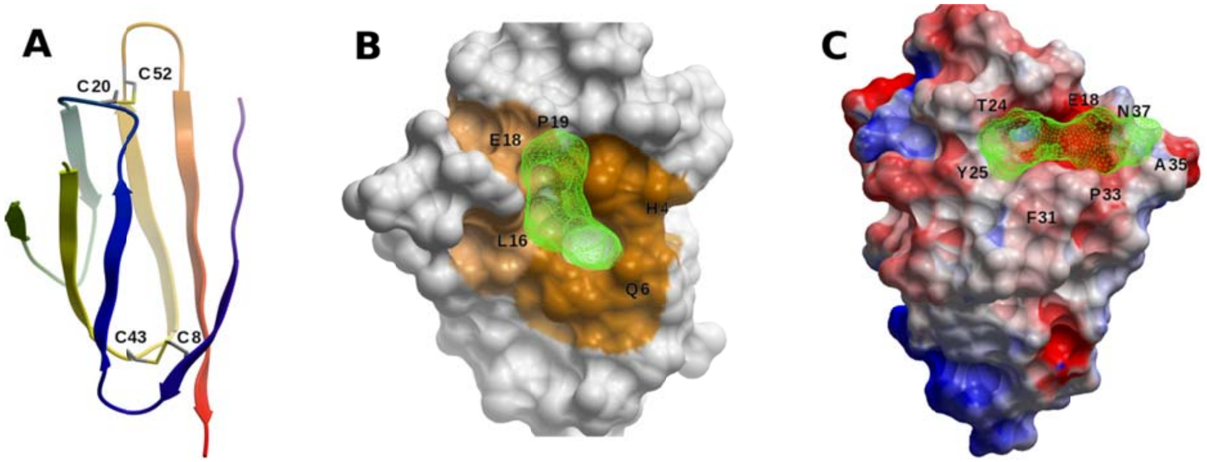


**Figure S14: Structure of SARS-CoV-2 orf7a.** (A) Ribbon representation of orf7a N-terminal ectodomain, colored the N-terminus (blue) to the C-terminus (red). The two disulfide bonds are shown. (B) A cryptic site identified in orf7a (colored by cryptic site score, light brown, intermediate; brown, high), with a neighboring consensus site (green mesh). (C) A hot-spot limited by amino acids E18, T24, Y25, F31, P33, A35, N37, and F50, shown as a green mesh. The surface of orf7a is colored according to the electrostatic potential (red, negatively charged; blue, positively charged, white, neutral).

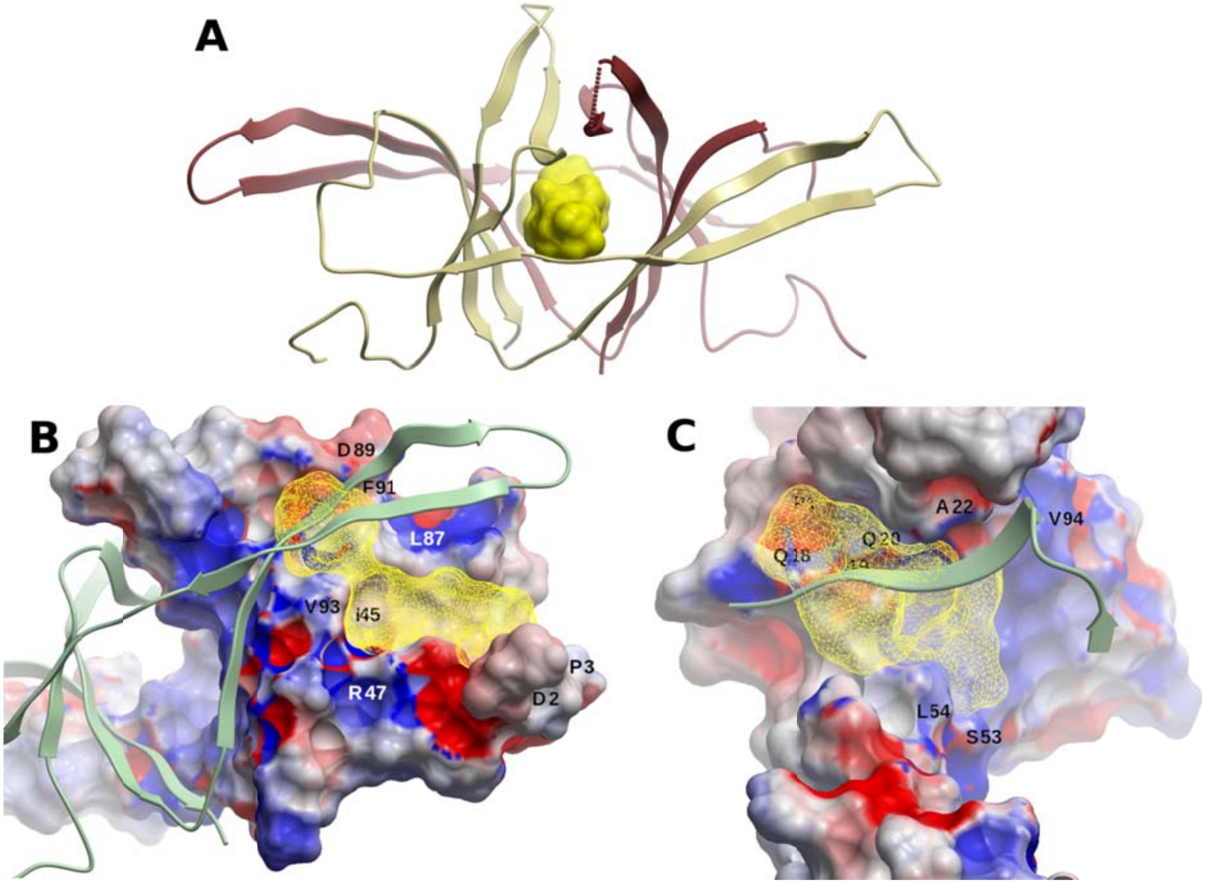


**Figure S15: The SARS-CoV-2 orf9b homodimer** (A) Structure of the orf9b homodimer. The hydrophobic central cavity is displayed in yellow. (B,C) Potential druggable sites in a orf9b monomer (yellow mesh representation). The surface of orf9b is colored according to the electrostatic potential (red, negatively charged; blue, positively charged, white, neutral). The second protomer is displayed as ribbon (in C, only a portion is shown, for the sake of clarity). A small molecule binding to these potentials sites might interfere with PPI between protomers, thus precluding homodimerization.
